# A Sentence Supramodal Areas Atlas (Sensaas) Based on Multiple Task-Induced Activation Mapping and Graph Analysis of Intrinsic Connectivity in 144 Healthy Right-Handers

**DOI:** 10.1101/273227

**Authors:** L Labache, M Joliot, J Saracco, G Jobard, I Hesling, L Zago, E Mellet, L Petit, F Crivello, B Mazoyer, N Tzourio-Mazoyer

**Affiliations:** Univ. Bordeaux, IMN, UMR 5293, F-33000 Bordeaux, France; CNRS, IMN, UMR 5293, F-33000 Bordeaux, France; CEA, GIN, IMN, UMR 5293, F-33000 Bordeaux, France; Univ. Bordeaux, IMB, UMR CNRS 5251, F-33405 Talence, France; INRIA Bordeaux Sud-Ouest, CQFD, INRIA, UMR 5251, F-33405 Talence, France

**Keywords:** fMRI, left hemisphere, language production, reading, comprehension, atlas, intrinsic connectivity, graph analysis, right-handers, resting-state

## Abstract

We herein propose an atlas of 32 sentence-related areas based on a 3-step method combining the analysis of activation and asymmetry during multiple language tasks with hierarchical clustering of resting-state connectivity and graph analyses. 144 healthy right-handers performed fMRI runs based on language production, reading and listening, both with sentences and lists of over-learned words. Sentence minus word-list BOLD contrast and left-minus-right BOLD asymmetry for each task were computed in pairs of homotopic regions of interest (hROIs) from the AICHA atlas. Thirty-two hROIs were identified that were conjointly activated and leftward asymmetrical in each of the 3 language contrasts. Analysis of resting- state temporal correlations of BOLD variations between these 32 hROIs allowed the segregation of a core network, SENT_CORE including 18 hROIs. Resting-state graph analysis applied to SENT_CORE hROIs revealed that the pars triangularis of the inferior frontal gyrus and the superior temporal sulcus were hubs based on their degree centrality, betweenness, and participation values, corresponding to epicentres of sentence processing. Positive correlations between DC and BOLD activation values for SENT_CORE hROIs were observed across individuals and across regions regardless of the task: the more a SENT_CORE area is connected at rest the stronger it is activated during sentence processing. DC measurements in SENT_CORE may thus be a valuable index for the evaluation of inter-individual variations in language areas functional activity in relation to anatomical or clinical patterns in large populations. SENSAAS (SENtence Supramodal Areas AtlaS), comprising the 32 supramodal sentence areas, including SENT-CORE network, can be downloaded at http://www.gin.cnrs.fr/en/tools/.

## Introduction

Defining language areas is a complex enterprise because of the numerous possible approaches currently available to identify language-related regions. The gold standard is to consider that language areas correspond to regions wherein lesions lead to aphasia. Even when limiting the definition of language areas to that of essential language areas, different identification methods exist that provide various kinds of information. Wada testing allows identification of the hemisphere controlling language but does not provide regional information (Wada and Rasmussen, 1960). By contrast, surgical cortical stimulation studies have documented left hemisphere language areas in large samples of patients (Ojemann et al., 1989, Tate et al., 2014) (Tate et al., 2014), but such mapping of eloquent areas is still limited to the cortical regions available to the neurosurgeon and is conducted in patients having potentially modified language organization. The probabilistic mapping of lesions combined with fine-grained aphasic patient evaluations of language performance have provided the community with very accurate descriptions of essential language areas (Dronkers and Ogar, 2004, Dronkers et al., 2004) although this very important approach does not reveal how these cortical areas are organized in networks. Because each multiple cortical area altered by a given pathology is not involved in the language deficit, the comprehensive identification of language areas from lesions is a complex issue (see (Genon et al., 2018) for a review).

Functional neuroimaging provides a way to map multiple areas activated during the completion of various language tasks in a large number of individuals. Furthermore, neuroimaging methodology is very efficient at compiling results obtained in multiple laboratories across the world, thereby allowing meta-analyses across laboratories that provide the location of areas activated at an acceptable spatial resolution within a common normalization space for a variety of language tasks. Similar to the results obtained with cortical stimulation (Ojemann et al., 1989), meta-analyses of neuroimaging data have provided the landscape of the left hemisphere cortical areas involved in language tasks in healthy individuals, which covers nearly the entire hemisphere surface (Price, 2000, Vigneau et al., 2006, Price, 2010, 2012).

Despite the vast amount of information obtained from the methods cited above, an atlas of left hemisphere language areas in healthy individuals having a typical left hemisphere dominance for language is still lacking, and with respect to language areas, the absence of a consensus is clear. The posterior part of the superior temporal gyrus and the supramarginal gyrus (Tomasi and Volkow, 2012, Klingbeil et al., 2017) are phonological regions that can be found under the label «Wernicke’s area», while lesion-based studies (Dronkers and Ogar, 2004) (Yourganov et al., 2015) as well as lesion studies in association with activation studies (Saur et al., 2006) have shown deep aphasia associated with lesions of the posterior region of the middle temporal gyrus and superior temporal sulcus (Binder, 2015, 2017). There is greater consistency concerning the location of frontal language areas under the label of Broca’s area because its original definition was anatomical. Most people define Broca’s area as the pars triangularis of the inferior frontal gyrus (Clos et al., 2013, Friederici and Gierhan, 2013, Yourganov et al., 2015). However, the extent of Broca’s area in the left frontal lobe varies, and the anterior insula (Baldo et al., 2011) is sometimes added, as reviewed in Amunts (Amunts and Zilles, 2012). Moreover, posterior lesions can also lead to Broca’s aphasia (Richardson et al., 2012), demonstrating that these anterior and posterior language poles work tightly together. This relation enhances the importance of networks in cognitive processing, as defined by Fuster (Fuster and Bressler, 2012). An atlas of language areas and networks in healthy individuals would thus be a useful tool, especially when individual task-induced mapping is not available. This atlas would be especially helpful for patients having difficulties completing language tasks and for the exploration of genetic language bases in large cohorts of individuals, in cohorts targeting normal or pathological brains, including those with developmental pathologies, and/or in individuals mapped for their anatomy and/or resting state while not performing a language task (Thompson et al., 2017).

To elaborate such an atlas, increasing the specificity for language areas that will be integrated is important because as uncovered by lesion studies, not all areas revealed by task-induced activation studies are essential language areas. Components of the task, such as monitoring, selecting, and holding the instructions, as well as paralinguistic processing, such as context, emotional and prosodic processing, are responsible for activations that exceed the essential language areas of the left hemisphere. The strong right hemisphere activations observed with functional imaging during various language tasks have even led some authors to claim that neuroimaging methods are not adequate to map language regions (Sidtis, 2007). One way to overcome this issue is using appropriate reference tasks. To discriminate language areas among those involved in the completion of a given task, Binder has suggested using well-designed reference tasks. The idea is to remove the non-specific or non-lateralized activations of primary areas and/or executive regions by applying the difference paradigm (Binder, 2011). Compared to a non-verbal reference, the use of a verbal reference tasks allows left hemisphere language areas to be specifically highlighted, as shown by Ferstl’s meta-analysis (Ferstl et al., 2007). The use of verbal reference tasks with functional magnetic resonance imaging (fMRI) has proven to successfully measure activation asymmetry, a proxy of language dominance strongly concordant with Wada testing. Note that this is true whether hemispheric or regional asymmetry of activations is used for the evaluation of language hemispheric dominance (review in (Dym et al., 2011)).

Thus, asymmetry represents an additional method for increasing the specificity of identifying left hemisphere language areas. Typical language organization, seen in 90% of the healthy population (Mazoyer et al., 2014) and 97% of healthy right-handers (Zago et al., 2017), is characterized by a strong left hemisphere dominance, giving rise to regional leftward asymmetries in fMRI. Adding to the detection of activated areas (by comparison to a high-level verbal reference task), a criterion based on leftward asymmetry would certainly increase the specificity of identifying left hemisphere language areas.

Another difficulty in identifying essential language areas with functional imaging is the fact that different tasks lead to different patterns of activation. One way to overcome this difficulty is to combine several language tasks in the same participant and apply conjunction analyses to unravel the activated and asymmetrical regions independent of the type of task or modality involved (Papathanassiou et al., 2000, Jobard et al., 2007, Dodoo-Schittko et al., 2012).

Finally, the task-induced approach does not provide any information on how the different activated areas are organized. The co-activation of a group of regions does not indicate that they are all strongly functionally connected and thus constitute a network. Resting-state intrinsic connectivity has proven to be capable of identifying the organization of brain network cognition. A good illustration of this concept is provided by Turken and Dronkers, who conducted correlation analysis in resting-state images of healthy participants using the posterior middle temporal gyrus region as the seed (Turken and Dronkers, 2011). In this work, this seed region was selected because its lesion was associated with strong comprehension deficits in aphasics, and its location was previously identified by probabilistic lesion mapping (Dronkers et al., 2004). Using this approach, Turken et al. revealed a network of areas connected at rest that support speech comprehension in healthy individuals. Investigating intrinsic connectivity would thus be an interesting means to investigate the networks existing at rest among the areas activated during language tasks. Connectivity measures can provide essential information on how regions are connected and how they are organized in networks. Graph analysis methodology applied on resting-state connectivity also permits the measurement of the connectivity strength of each region with all other regions of a given network to which it belongs, thereby characterizing its role in the network. In particular, identifying the topological roles of the regions is possible, i.e., identifying hubs, regions essential to a given network and therefore essential to the cognitive function(s) they support (Sporns et al., 2007).

To propose an atlas of left hemisphere high-order language areas, we first combined multiple language fMRI task-induced activation mapping and conjunction analysis to select a set of both activated and leftward asymmetrical areas during sentence processing. Second, we clustered the regions identified in the first step into networks based on their intrinsic connectivities at rest. Third, we applied graph analysis to characterize the roles of the regions in communication within and across networks. To this end, we utilized BIL&GIN, a database dedicated to the study of hemispheric specialization (Mazoyer et al., 2015), and selected 144 right-handers who were mapped during sentence production, reading and listening tasks compared to the production, reading and listening of lists of words, respectively. All but 6 participants were also mapped during the resting state. Most investigations of the resting-state and task-induced activation networks have relied on whole-brain comparisons between the functional connectivities measured in these two conditions (review in Wig (Wig, 2017)), although they correspond to very different physiological states (Raichle and Mintun, 2006, Raichle, 2015). Here, we aimed to find resting-state markers of left hemisphere activation in discrete language areas to provide a comprehensive tool for further research on the interindividual variability of language areas. Such markers of language activation are likely to be of interest for studies in which no task-induced activations are documented but instead include a resting-state acquisition. Consequently, we used homotopical regions of interest (hROIs) from the AICHA atlas, a functional atlas obtained from intrinsic connectivity analysis (Joliot et al., 2015). We used AICHA hROIs because (1) we needed an atlas suitable for functional imaging and, in that respect, AICHA, which was elaborated from resting-state connectivity, is optimal for analysing functional data; (2) AICHA has been specifically designed to identify functionally homotopic regions of interest, enabling the accurate computation of functional asymmetries since it avoids the potential bias that anatomical and functional areas do not strictly overlap. The different sets of hROIs corresponding to the supramodal sentence processing areas of the proposed atlas (SEN-SAAS) are available in the Montreal Neurological Institute (MNI) space at http://www.gin.cnrs.fr/en/tools/.

## Material and methods

### Participants

From the BIL&GIN database, we selected 144 healthy right-handers (72 women) who completed the fMRI battery, including several language tasks (Mazoyer et al., 2015). The sample mean age was 27 years (SD = 6 years), and the women were two years younger than the men (women 26 ± 5; men: 28 ± 7, p = 0.053). The mean educational level of the participants was 16 years (SD = 6 years), with no significant difference between the men and women (p > 0.05). All participants reported themselves as right-handed; their mean normalized finger tapping test asymmetry ([(right number of taps - left number of taps) / (left + right number of taps)] x 100) was 6.25 (SD = 4.3), and their mean Edinburgh score was 93.5 (SD = 11), confirming their right-handedness. There was no difference between gender for the Edinburgh score (p = 0.47), although there was a slightly stronger rightward manual laterality in women (finger tapping test asymmetry in women: 6.9 ± 3.8; men: 5.7 ± 4.7, p = 0.08, controlling for age).

Of these participants, 138 (mean age 27 years (SD = 6 years), 68 women) also completed a resting-state fMRI (rs-fMRI) acquisition lasting 8 minutes. Note that this resting-state acquisition was performed on average 9 months (SD = 9.6 months) before the language task acquisition in all but 5 cases. In these 5 cases the resting-state acquisition occurred approximately one year after the language session (range [11.2 - 13.8] months).

### Image acquisition and processing

#### Structural imaging

Structural images were acquired using the same 3T Philips Intera Achieva scanner including high-resolution T1-weighted volumes (sequence parameters: TR, 20 ms; TE, 4.6 ms; flip angle, 10°; inversion time, 800 ms; turbo field echo factor, 65; sense factor, 2; field of view, 256 × 256 × 180 mm3; isotropic voxel size 1 × 1 × 1 mm3). For each participant, the line between the anterior and posterior commissures was identified on a mid-sagittal section, and the T1-MRI volume was acquired after orienting the brain in the bi-commissural coordinate system. T2*-weighted multi-slice images were also acquired (T2*-weighted fast field echo (T2*-FFE), sequence parameters: TR = 3,500 ms; TE = 35 ms; flip angle = 90°; sense factor = 2; 70 axial slices; 2 x 2 x 2 mm3 isotropic voxel size).

#### Task-induced functional imaging

##### Training

To ensure proper task execution, the participants were trained outside the scanner in the hour preceding the fMRI session. The training used stimuli that were of the same nature but different from those used during the fMRI session.

##### Language tasks

Three runs were administered to the participants. They included a sentence task involving phonological, semantic, prosodic and syntactic processing and a word-list reference task, a less complex, albeit high-level, verbal task. To achieve homogeneity in the sentence task material, 51 line drawings illustrating the stories of ‘Le petit Nicolas’ (Little Nicholas), a classic French children’s series, were used. The three tasks consisted of a randomized alternation of event-related trials devoted to sentence processing, with event-related trials devoted to the verbal reference task, i.e., lists of words. The drawings used for the reference task were scrambled versions of the line drawings, and the stimuli presented either orally or visually were lists of months, days and/or seasons. Within each trial, the subject was shown either a line drawing or a scrambled drawing for 1 s, immediately followed by a central fixation crosshair. While fixating the cross, the subject performed either the sentence task or the word reference task. Once the task was completed, a low-level reference task, detecting the transformation of a centrally displayed cross into a square, was presented. When the subjects detected this change, they were asked to press a button with their index finger of the assigned hand. The square was then displayed until the end of the trial. This second part of the trial, which lasted at least half of the total trial duration, aimed at refocusing the subject’s attention to a non-verbal stimulus and controlling for manual motor response activation, which was also present in the first part of the trial. A 12-s presentation of a fixation crosshair preceded and followed the first and last trial. Note that except during the drawings display, the subjects were asked to keep fixating the cross, and the star and square were then presented on the centre of the screen.

###### Sentence and list of word production tasks

During the production run, after seeing a Little Nicholas line drawing, the subject was instructed to covertly generate a sentence beginning with a subject (The little Nicholas…, The gentleman…) and a complement (with his satchel…, in shorts…, with glasses…), followed by a verb describing the action taking place and ending with an additional complement of a place (in the street…, in the playground…, on the beach…) or a manner (with happiness…, nastily…). When a scrambled drawing was displayed, the subject was asked to covertly generate the list of the months of the year. The production paradigm randomly alternated ten 18-s trials of sentence generation with ten 18-s trials of generating the list of months. The response time limit, indicated by the transformation of the cross in a star, was 9 s, including the 1-s drawing display. The entire experimental run lasted 6 min and 24 s. The mean sentence production time was 5,617 ms (SD = 935 ms), while the mean duration of wordlist production was 5,249 ms (SD = 1,131 ms).

###### Sentence and list of word listening tasks

When a Little Nicholas line drawing was displayed, the subject was instructed to carefully listen to a sentence dealing with the line drawing and click at the end of the sentence. For the LISN, when a scrambled drawing was displayed, he/she was instructed to listen to the list of the months, days of the week and/or seasons and click at the end of the list.

The paradigm consisted of a randomized alternation of thirteen 14-s sentence listening trials with thirteen 14-s list listening trials. The mean durations of auditory presentation were 4,371 ± 468 ms for the sentences and 4,386 ± 484 ms for the lists. The entire experimental run lasted 6 min and 28 s. The reaction times after sentence and list listening were 387 ms (SD = 125 ms) and 478 ms (SD = 97 ms), respectively.

###### Sentence and list of word reading tasks

Like in the other two tasks, when a line drawing was displayed, the subject was instructed to read a sentence based on the line drawing. When a scrambled drawing was displayed, he/ she was instructed to read the list of months, days of the week and/or seasons.

The paradigm consisted of a randomized alternation of thirteen 14-s sentence reading trials with thirteen 14-s list reading trials. The entire experimental run lasted 6 min and 28 s. The average time for reading sentences was 3,729 ms (SD = 567 ms), while reading the lists of words required 4,412 ms (SD = 602 ms).

##### Debriefing the fMRI tasks

Right after the fMRI sessions, the participants were asked to rate the difficulty of the task on a 5-point scale (1-easy to 5-very difficult) and answer some debriefing questions about how they accomplished the task.

The production task had the highest task difficulty score reported by the participants (2.73), while the reading and listening tasks had low scores (1.14 and 1.20, respectively). All participants were able to recollect the sentence they produced when presented with the corresponding drawing for at least 5 of 10 images (mean: 9.43, SD: 0.96), with the mean number of words per sentence being 12.4 (SD = 2).

#### Functional image acquisition

The functional volumes were acquired as T2*-weighted echo-planar EPI images (TR = 2 s; TE = 35 ms; flip angle = 80°; 31 axial slices with a 240 x 240 mm^2^ field of view and 3.75 x 3.75 x 3.75 mm^3^ isotropic voxel size). In the three runs, 192, 194 and 194 T2*-weighted volumes were acquired for the production, listening and reading sentence tasks, respectively.

#### Resting-state functional imaging (rs-fMRI)

Spontaneous brain activity was monitored for 8 minutes (240 volumes) using the same imaging sequence (T2*-weighted echo-planar images) as that used for the language tasks.

Immediately prior to rs-fMRI scanning, the participants were instructed to “keep their eyes closed, to relax, to refrain from moving, to stay awake and to let their thoughts come and go”.

#### Image analysis

##### Functional imaging analysis common to task-induced and resting-state acquisitions

For each participant, (1) the T2*-FFE volume was rigidly registered to the T1-MRI; (2) the T1-MRI was segmented into three brain tissue classes (grey matter, white matter, and cerebrospinal fluid; and (3) the T1-MRI scans were normalized to the BIL&GIN template including 301 volunteers from the BIL&GIN database (aligned to the MNI space) using the SPM12 “normalise” procedure with otherwise default parameters.

For each of the 3 fMRI runs, data were corrected for slice timing differences. To correct for subject motion during the runs, all the T2*-weighted volumes were realigned using a 6-parameter rigid-body registration. The EPI-BOLD scans were then registered rigidly to the structural T2*-FFE image. The combination of all registration matrices allowed for warping the EPI-BOLD functional scans to the standard space with a single trilinear interpolation.

##### Specific task-induced functional imaging analysis

First, a 6-mm full width at half maximum (Gaussian filter was applied to each run. Global linear modelling (statistical parametric mapping (SPM), http://www.fil.ion.ucl.ac.uk/spm/) was used for processing the task-related fMRI data. For each participant, BOLD variations corresponding to each sentence versus the list belonging to the same run were computed (sentence minus word-list production (PROD _SENT-WORD_), sentence minus word-list reading (READ _SENT-WORD_), and sentence minus word-list listening (LISN_SENT-WORD_)). Finally, contrast maps (defined at the voxel level) were subjected to ROI analysis. BOLD signal variations were measured in 192 pairs of functionally defined hROIs of the AICHA atlas (Joliot et al., 2015) adapted to SPM12, excluding 7 hROI pairs belonging to the orbital and inferior-temporal parts of the brain in which signals were reduced due to susceptibility artefacts. For each participant, we computed contrast maps of the 3 language conditions. We then calculated the right and left hROI BOLD signal variations for each of the 185 remaining pairs by averaging the contrast BOLD values of all voxels located within the hROI volume.

##### Specific analysis of resting-state functional images

Time series white matter and cerebrospinal fluid (individual average time series of voxels that belonged to each tissue class) and temporal linear trends were removed from the rs-fMRI data series using regression analysis. Additionally, rs-fMRI data were temporally filtered using a least squares linear-phase finite impulse response filter design bandpass (0.01 Hz – 0.1 Hz). For each participant and hROI (the same 185 homotopic ROIs as those used in the task- induced analysis), an individual BOLD rs-fMRI time series was computed by averaging the BOLD fMRI time series of all voxels located within the hROI volume.

## PART 1. Identification and Characterization of HROIs Exhibiting Both Leftward Activation and Leftward Asymmetrical Activation in All 3 Tasks

To complete the identification of high-order language areas, we first searched for hROIs that were both significantly co-activated and significantly leftward asymmetrical on average among the 144 participants during the PROD _SENT-WORD_, READ_SENT-WORD_, and LISN_SENT-WORD_ tasks.

## Statistical analysis

### hROI selection

Using JMP13 (www.jmp.com, SAS Institute Inc., 2012), conjunction analysis was conducted to select the left-hemisphere hROIs exhibiting BOLD signal variations that were both significantly positive and significantly larger than that in their right counterparts in all 3 tasks. An hROI was selected whenever it was significantly activated in each of the 3 task contrasts using a significance threshold set to p < 0.05 per contrast. The significance threshold for the conjunction of activation in the 3 tasks was thus 0.05 x 0.05 x 0.05 = 1.25×10^−4^. The second criterion for hROI selection was the existence of a significant leftward asymmetry in each of the 3 task contrasts, the threshold of significance of this second conjunction being again 1.25×10^−4^. Finally, since to be selected, a given hROI had to fulfil both criteria, the overall significance threshold for the conjunction of conjunction analyses was 1.5×10^−8^ = (1.25×10^−4^)2.

## Results

### hROI selection

Among the 80 hROIs jointly activated in the 3 contrasts (Table 1), 46 also showed joint asymmetries. In total, 32 hROIs showed both joint activation on the left and joint asymmetry (Figure 1, Table 2).

**Fig. 1.**
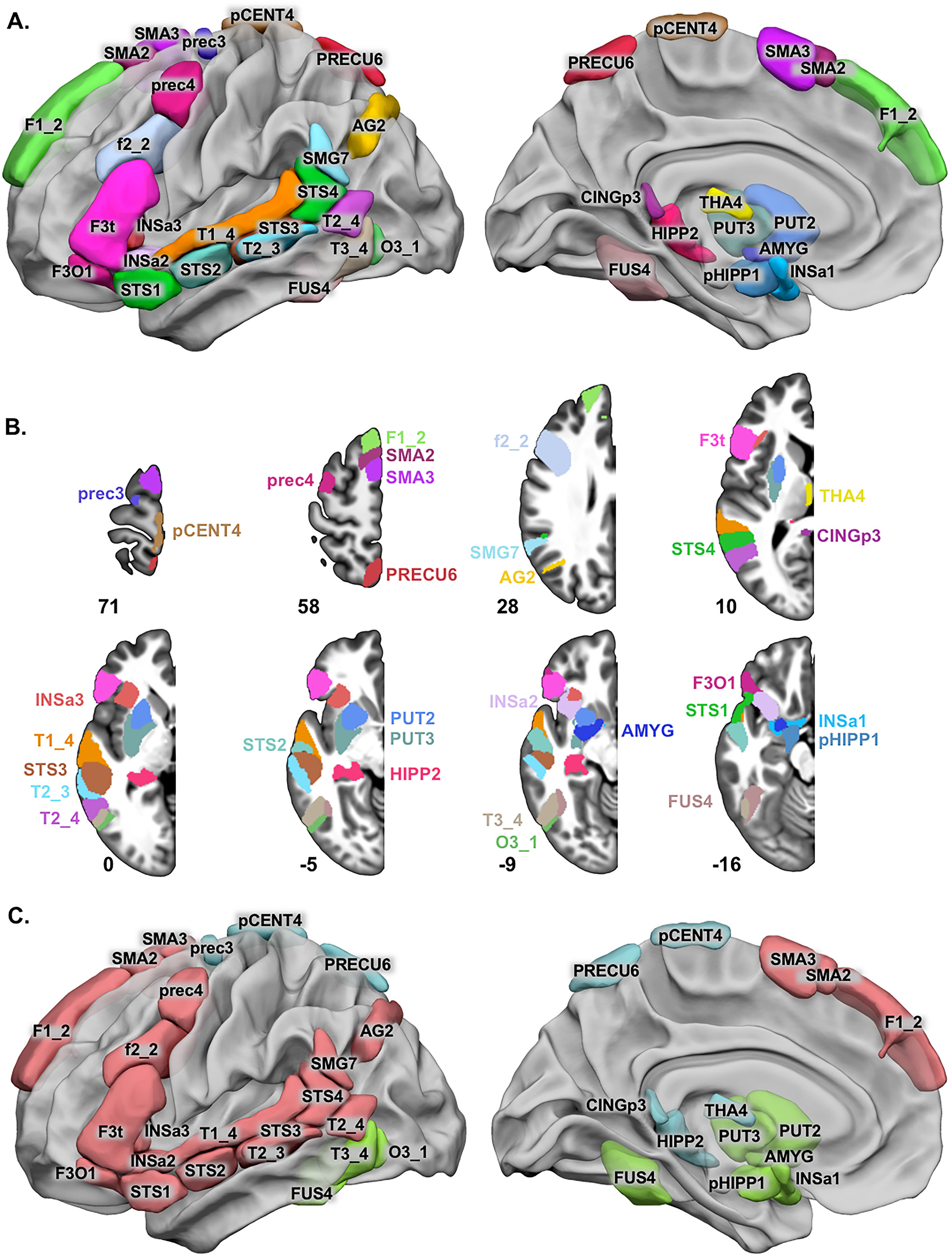
Locations of the 32 hROIs co-leftward activated and co-leftward asymmetrical during the completion of 3 sentence minus word-list tasks by 144 healthy right-handers and corresponding networks after hROI clustering based on resting-state connectivity. **a** Left lateral view of 3D surfaces rendering the 32 hROIs on the BIL&GIN display template in the MNI space with Surf Ice (https://www.nitrc.org/projects/surfice/) software. **b** Representation of hROIs on left hemisphere axial slices from the BIL&GIN display template; the hROI numbers correspond to the z-axis in the MNI space. **c** Lateral and medial views of the 4 identified networks. SENT_CORE network: pink, SENT_MEM: light blue and SENT_VISU: green. Correspondences between the abbreviations and the full names of the AICHA atlas can be found in Table 2.

**Table 1.** Results of conjunction analyses across each sentence minus word-list contrasts for production (PROD_SENT-WORD_), listening (LISN_SENT-WORD_) and reading (READ_SENT-WORD_) tasks in terms of the number of hROIs. Numbers of hROIs with significant left activation, leftward asymmetry or conjunction of activation and asymmetry for the 3 “sentence minus word” contrasts.

|  | L<br>activation | L<br>asymmetry | Conjunction of activation<br>and asymmetry |
| --- | --- | --- | --- |
| $PROD_{SENT-WORD}$ | 133 | 93 | 75 |
| $LISN_{SENT-WORD}$ | 116 | 73 | 64 |
| $READ_{SENT-WORD}$ | 97 | 60 | 43 |
| Conjunctions<br>of 3 contrasts | 80 | 46 | 32 |

**Table 2.**
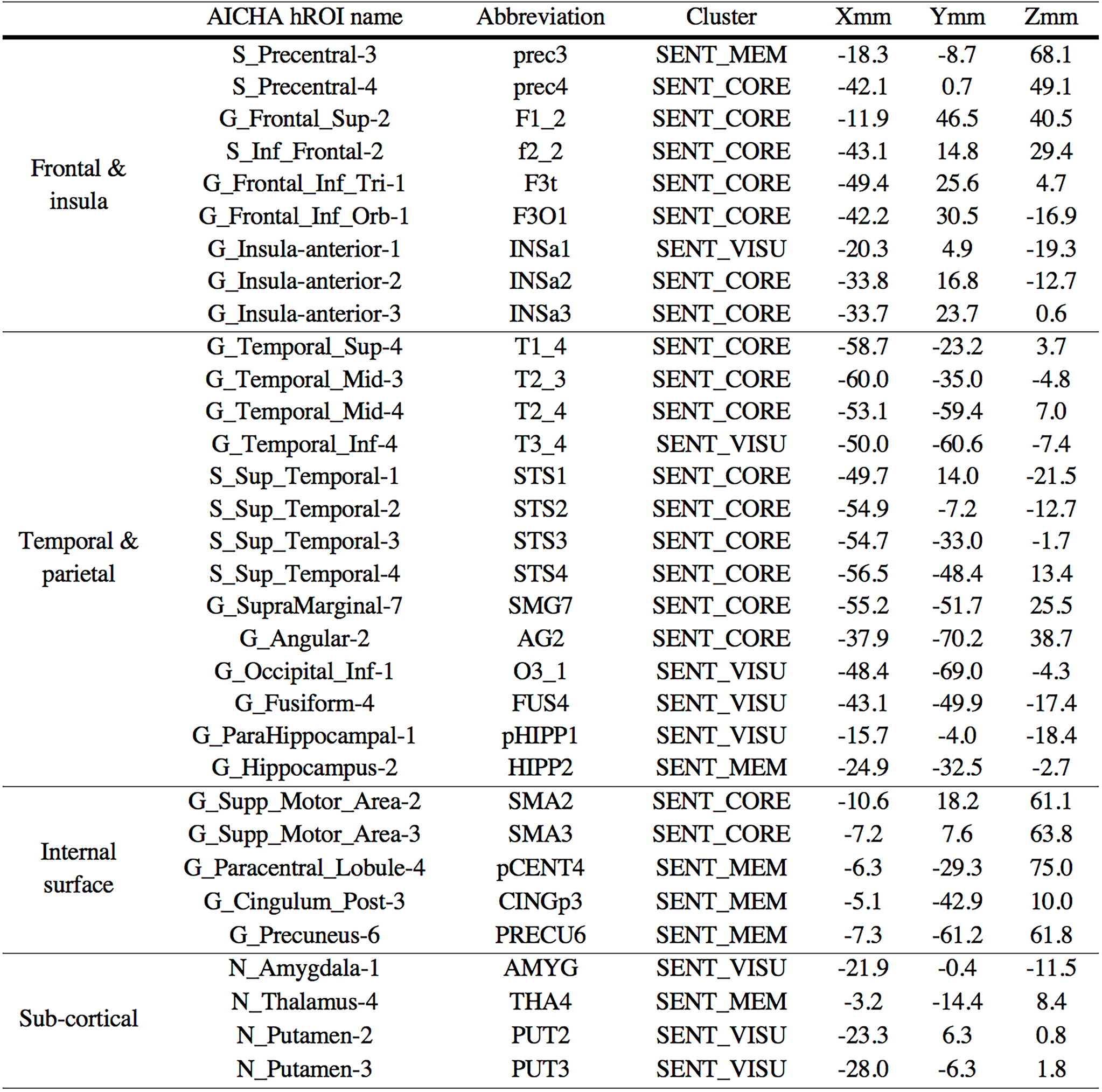
Names and abbreviations of the 32 hROIs showing joint left activation and left asymmetry during the three sentences minus word-list contrasts for production (PROD_SENT-WORD_), listening (LISN_SENT-WORD_) and reading (READ_SENT-WORD_) tasks; the network label to which they were clustered; and their coordinates in MNI space after SPM12 normalization of the AICHA atlas.

On the lateral surface of the left frontal lobe, the regions having both joint activation and leftward asymmetry during the 3 language tasks covered the left inferior frontal gyrus (pars triangularis: F3t and pars opercularis: F3O1), the adjacent inferior frontal sulcus (f2_2), the junction of the middle frontal gyrus with the precentral sulcus (prec4), and the upper part of the precentral sulcus (prec3) located dorsally to prec4. The medial part of the superior frontal gyrus (F1_2), the upper paracentral gyrus (pCENT4), and the pre-superior motor areas (SMA2 and SMA3) were also part of these areas in the medial frontal lobe. Two hROIs were located within the anterior insula (INSa2 and INSa3), while the INSa1 hROI was located medially and ventrally close to the amygdala. On the lateral surface of the temporal lobe, the hROIs overlapped the entire length of the superior temporal sulcus (STS2, STS3 and STS4), extending to the temporal pole anteriorly (STS1), to the superior temporal gyrus dorsally (T1_4), to the supramarginal (SMG7) and angular gyri (AG2) posteriorly, crossing the middle temporal gyrus (T2_3 and T2_4) and joining the inferior temporal gyrus (T3_4), the inferior occipital gyrus (O3_1), and ventrally the fusiform gyrus (FUS4). Regions located within the hippocampus (HIPP2), parahippocampal gyrus (pHIPP1) and amygdala (AMYG) were also part of the selected areas. In the posterior medial wall, the dorsal part of the precuneus (PRECU6) together with the posterior cingulum (CINGp3) was selected using this approach. Sub-cortical areas jointly activated and leftward asymmetrical during the 3 tasks covered almost the entire putamen (PUT2 and PUT3_4) and a thalamic hROI located medially (THA4).

## PART 2. Identification of Networks Based ON the Resting-State Connectivity Matrix of the 32 HROIs Co-Activated and Co-Leftward Asymmetrical During the 3 Sentence Minus Word-List Tasks

In a second step, we investigated the intrinsic functional organization of the 32 hROIs selected in the first step. We computed the intrinsic connectivity matrix between these 32 hROIs for the subsample of 138 right-handed participants who completed a resting-state acquisition. We completed a hierarchical clustering analysis of this intrinsic connectivity matrix to identify temporally coherent networks within this set of hROIs.

## Methods

### Calculation of the intrinsic connectivity matrix

An intrinsic connectivity matrix was calculated for each of the 138 individuals and for each of the 496 possible pairs of hROIs (N x (N-1))/2, with N = 32). The intrinsic connectivity matrix off-diagonal elements were the Pearson correlation coefficients between the rs-fMRI time series of the hROI pairs. The intrinsic connectivity matrix diagonal elements were set to zero because no information on the correlation for a specific hROI with itself exists (Rubinov and Sporns, 2010). The individual intrinsic connectivity matrix was then Fisher z-transformed before being averaged over the subsample of 138 individuals, thereby producing a mean intrinsic connectivity matrix.

### Identification and characterization of networks

The agglomerative hierarchical cluster analysis (aHCA) method was applied to extract brain networks from this mean intrinsic connectivity matrix. We first transformed the Pearson correlation (r_i,j_) between hROI i and hROI j into a distance (d_i,j_) using the equation d_i,j_ = (1- r_i,j_)/2, as in Doucet (Doucet et al., 2011), resulting in a 32×32 dissimilarity matrix. According to Lance and Williams (Lance and Williams, 1979), the previous equation is “unequivocally the best” to transform a correlation into a distance on real data sets. Finally, we used an agglomerative hierarchical clustering algorithm (aHCA, (Sneath and R, 1973)) for clustering and the Ward distance (Ward Jr, 1963) for aggregating the different hROIs into clusters. The number of clusters (networks) was determined using the R library NbClust (Charrad et al., 2014). This package provides 30 statistical indices for determining the optimal number of clusters and proposes the best clustering scheme from the different results obtained by varying all combinations of number of clusters for the chosen method, in this case, aHCA with Ward’ s distance. We chose the number of clusters that fulfilled a maximum of indices.

To characterize each network, we calculated its mean volume activation for each task contrast as the sum of the activations of all hROIs composing the network weighted by their individual volume and then divided by the sum of their volumes. The same computation was performed for the right hemisphere equivalent of each network, which was then used for computing the left-minus-right asymmetry of each network activation. We then compared activation amplitude and asymmetry values across networks and across tasks using a mixed-model ANOVA.

### Reliability of the network identification across individuals

We used multiscale bootstrap resampling (Efron and Halloran, 1996) to assess the reliability of the identification of each cluster. In total, 10,000 multiscale bootstrap resampling datasets, including 50% to 140% of sample data from the 138 participants (Suzuki and Shimodaira, 2004), were processed. Applying the R package “pvclust” (Suzuki and Shimodaira, 2006) function to the multiscale bootstrap resampling outputs, we measured the approximately unbiased (AU) p-value for each cluster. The AU p-value for a network, the probability of this network occurring among the 138 participants, indicates the network’s reliability.

### Robustness of the network identification with respect to the clustering method

We also assessed the robustness of the clustering method by comparing its output to those of 3 other clustering methods: aHCA with the average distance method (instead of Ward’s), Gaussian mixture model, and k-means (see supplementary material, Table 1). Gaussian mixture modelling was conducted with the “Rmixmod” package with Normalized Entropy Criterion in order to find well-separated clusters and with a Gaussian model with diagonal variance matrices (Lebret et al., 2014).

We then compared the 4 different partitions through the adjusted Rand index (L and Arabie, 1985) allowing to get a similarity measure between 2 different classifications, an adjusted Rand index of 1 indicating that similar partitions.

### Temporal correlation across networks and significance

To compute the mean intrinsic functional correlations between two networks, we used the same methodology as that used to compute the mean intrinsic connectivity matrix (see above). First, for each individual and for each network, we computed the corresponding rs-fMRI time series by averaging the individual resting time series of all voxels of all hROIs belonging to this network.

Then, for each individual, we computed the Pearson correlation coefficients between all pairs of networks that we further Fisher z-transformed. Finally, each of these z-transformed coefficients was averaged across the sample of 138 individuals, providing a mean intrinsic functional correlation (r) for each pair of networks. We assessed the significance of each of these mean intrinsic functional correlations compared to 0 using a non-parametric sign test at the 0.05 significance level (Bonferroni correction for the number of network pairs).

## Results

### Identification and characterization of networks

Hierarchical clustering analysis revealed three networks from the selected set of 32 hROIs (Figure 2).

**Fig. 2.**
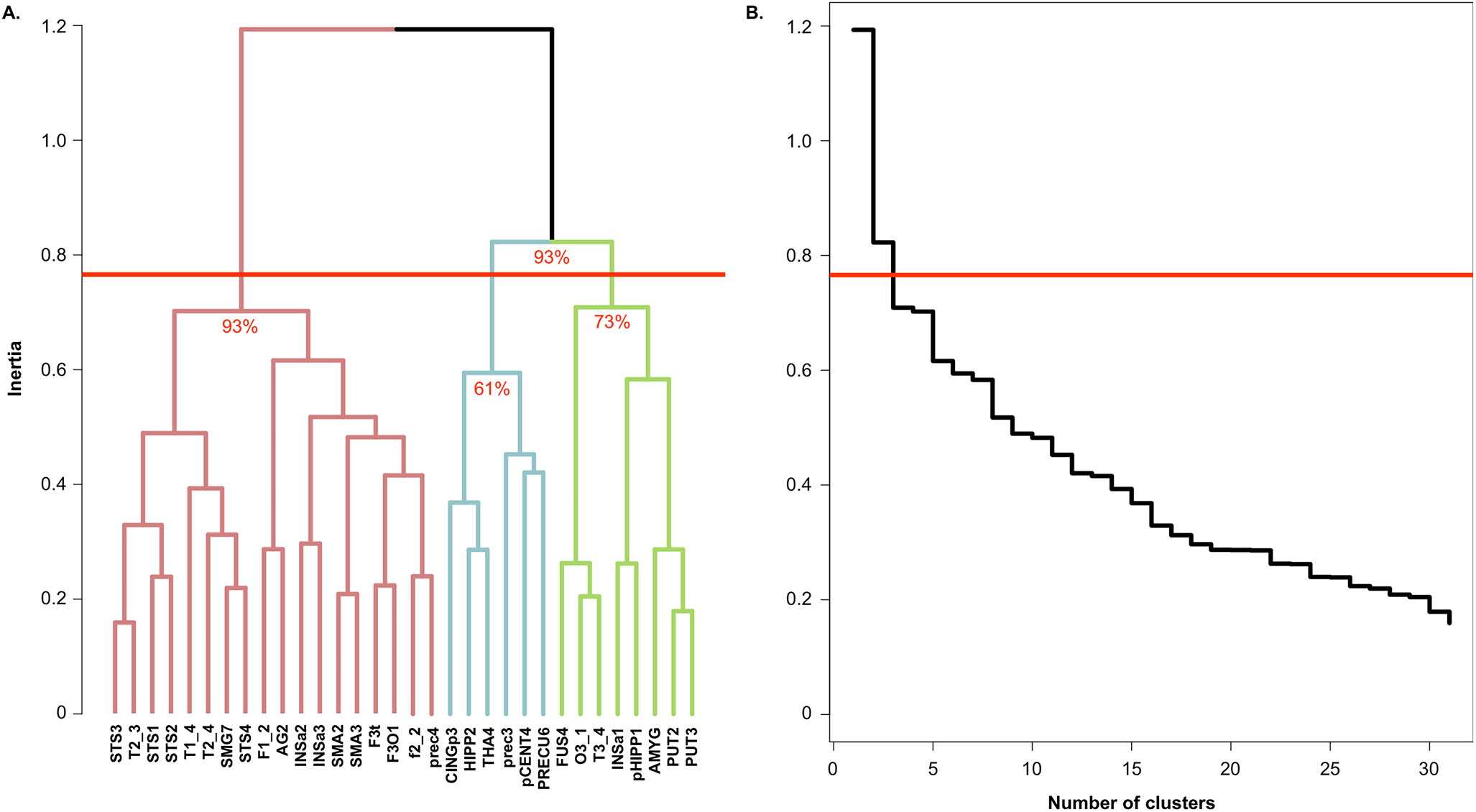
Results of the agglomerative hierarchical cluster analysis method. **a** Dendrogram of aHCA of the mean intrinsic connectivity matrix (SENT_CORE network: pink, SENT_MEM: light blue, SENT_VISU: green). Approximately Unbiased p-value are indicated for each identified network. **b** Scree plot of aHCA of the mean intrinsic connectivity matrix. For both graphs, the red horizontal line corresponds to the threshold applied to select the number of networks.

#### SENT_CORE network

The first network (pink in Figure 1C), termed SENT_ CORE, was composed of 18 hROIs and was the most distant from the 2 others in terms of inertia (pink in Figure 2A). SENT_CORE included all lateral and medial hROIs of the frontal lobe, apart prec3, pCENT4 and anterior insula INSa2 and INSa3. SENT_CORE also included all temporal and parietal hROIs of the lateral surface, except T3_4, which was aggregated with the network gathering visual hROIs.

We named this network SENT_CORE because it included essential sentence processing regions, as further described below. SENT_CORE was the largest network in terms of volume, as it was 9.2 times larger than SENT_ MEM and 3.4 times larger than SENT_VISU (Table 3).

**Table 3.** Mean volumetric activation (and standard deviation) of the 4 language networks in each sentence minus word-list contrast for production (PRODSENT-WORD), listening (LISNSENT-WORD) and reading (READSENT-WORD) tasks in 144 healthy right-handers. The mean volumetric activation for a network was calculated from the sum of the activations of the ROIs comprising the network weighted by their individual volumes and then divided by the volume of the cluster.

|  | Volume<br>(mm<sup>3</sup>) | Left activation |  |  | Leftward asymmetry |  |  |
| --- | --- | --- | --- | --- | --- | --- | --- |
|  |  | PROD<sub>SENT-WORD</sub> | LISN<sub>SENT-WORD</sub> | READ<sub>SENT-WORD</sub> | PROD<sub>SENT-WORD</sub> | LISN<sub>SENT-WORD</sub> | READ<sub>SENT-WORD</sub> |
| SENT_CORE | 83232 | 0.73 ± 0.31 | 0.43 ± 0.20 | 0.55 ± 0.26 | 0.41 ± 0.22 | 0.25 ± 0.14 | 0.28 ± 0.19 |
| SENT_MEM | 9024 | 0.40 ± 0.36 | 0.28 ± 0.23 | 0.21 ± 0.26 | 0.11 ± 0.17 | 0.08 ± 0.12 | 0.07 ± 0.14 |
| SENT_VISU | 24368 | 0.37 ± 0.27 | 0.30 ± 0.17 | 0.25 ± 0.21 | 0.16 ± 0.13 | 0.11 ± 0.08 | 0.12 ± 0.10 |

#### SENT_VISU network

This group of clusterized areas included the inferior temporal and occipital gyri laterally (T3_4, O3_1); the mid-fusiform (FUS4) ventrally; the parahippocampal region; the amygdala (AMYG1) and INSa1, close to the amygdala medially (Figure 1C, green); and the two hROIs of the putamen (PUT2 and PUT3). We labelled it SENT_VISU because it aggregated four hROIs acknowledged as involved in visual processing.

#### SENT_MEM network

This network (Figure 1C, light blue) included three regions of the medial wall —the paracentral gyrus (pCENT4), the precuneus (PRECU6) and the posterior cingulate (CINGp3)— and the posterior part of the hippocampus (HIPP2) as well as one frontal area at the upper end of the precentral sulcus (prec3) and the THA4 hROI located medially in the thalamus. We named it SENT_MEM because these posterior areas belong to both the posterior regions of the DMN involved in the posterior hippocampus in episodic memory, as further discussed below.

#### Profile comparisons between networks

In terms of mean voluminal activity (Table 3), ANOVA revealed a significant network effect, a task effect and a network x task interaction (all post hoc Tukey’s HSD: p < 10^−4^).

The interaction occurred because while in SENT_ CORE, there was greater activation in PROD, followed by READ and then by LISN (all p < 10^−4^), SENT_MEM and SENT_VISU had different profiles. Although the greatest activation values were also observed during PROD (all p < 0.003), activation values were significantly higher during LISN than during READ in both networks (SENT_VISU: p = 0.01; SENT_MEM: p = 0.0067).

ANOVA on asymmetries also revealed a significant network effect, a task effect and a network x task interaction (all post hoc Tukey’s HSD: p < 10^−4^). The interaction was due to different profiles of asymmetry in SENT_CORE than in the two other networks. In SENT_ CORE, the profile of asymmetries was the same as that of activation: larger in PROD, followed by READ and then by LISN (all post hoc Tukey’s HSD: p < 10^−4^). In SENT_ VISU, the asymmetry during PROD was slightly larger than that during READ (SENT_VISU: p = 0.0005; SENT_ MEM: p = 0.01) and larger than that during LISN (p = 0.0001). In addition, there was no difference in asymmetry between PROD and LISN in SENT_MEM (p = 0.075) and no difference in asymmetry between READ and LISN in SENT_VISU (p = 0.52) or SENT_MEM (p = 0.25). The task main effect was related to larger asymmetries in PROD and the network main effect to larger asymmetries in SENT_CORE (all p < 0.0001).

### Assessing the reliability of the identification of 2 networks across individuals in the 138 participants

The AU p-values provided by the multiscale bootstrap resampling method showed that each network of the first partition was reliable at levels of 93%, corresponding to SENT_CORE on one side and to the SENT_VISU and SENT_MEM on the other side (Figure 2). However, for the second partition, SENT_VISU and SENT_MEM were reliable at 73% and 61%, respectively, indicating a lower reliability.

### Robustness of the identified networks with respect to the clustering method

The SENT_CORE network was identified by all 4 clustering methods, including at least 13 of the 18 hROIs initially found with the aHCA method using Ward’s distance (see supplementary Table 1). The aHCA method using the average distance metric led to an adjusted Rand index of 1, indicating a clustering similar to that achieved using the aHCA method with Ward’s distance. Comparing the Gaussian Mixture Model and aHCA methods led to an adjusted Rand index of 0.76, indicating two highly similar partitions. Comparing the aHCA and k-means methods led to the lowest adjusted Rand index of 0.43.

Only one hROI of SENT_CORE (AG2) was segregated in SENT_MEM by Gaussian mixture modelling and by k-means, while all 17 other hROIs were classified together by all clustering methods but k-means (supplementary Table 1), although PUT3 joined SENT_CORE according to Gaussian mixture modelling. k-means classified INSa3, T2-4, T1-4 and f2-2 in SENT_VISU rather than in SENT_CORE.

Only prec3 was classified in SENT_VISU rather than SENT_MEM with both Gaussian mixture modelling and k-means, while pHIPP1 shifted from SENT_VISU to SENT_MEM only with k-means.

### Temporal correlation across networks and significance

The chord diagram shown in Figure 3 describes the average correlations between each pair of hROIs in the 3 networks. Strong and highly significant negative mean intrinsic correlations were found between SENT_CORE and SENT_MEM (R = −0.27; 92.03% of the participants showed a negative correlation, p < 10^−4^), and a positive correlation was present between SENT_MEM and SENT_ VISU (R = 0.058, 62.32% of the participants showed a positive correlation, p = 0.0024), while there was no significant correlation between SENT_CORE and SENT_VISU (R = −0.037; 56.52% of the participants showed a positive correlation, p = 0.074).

**Fig. 3.**
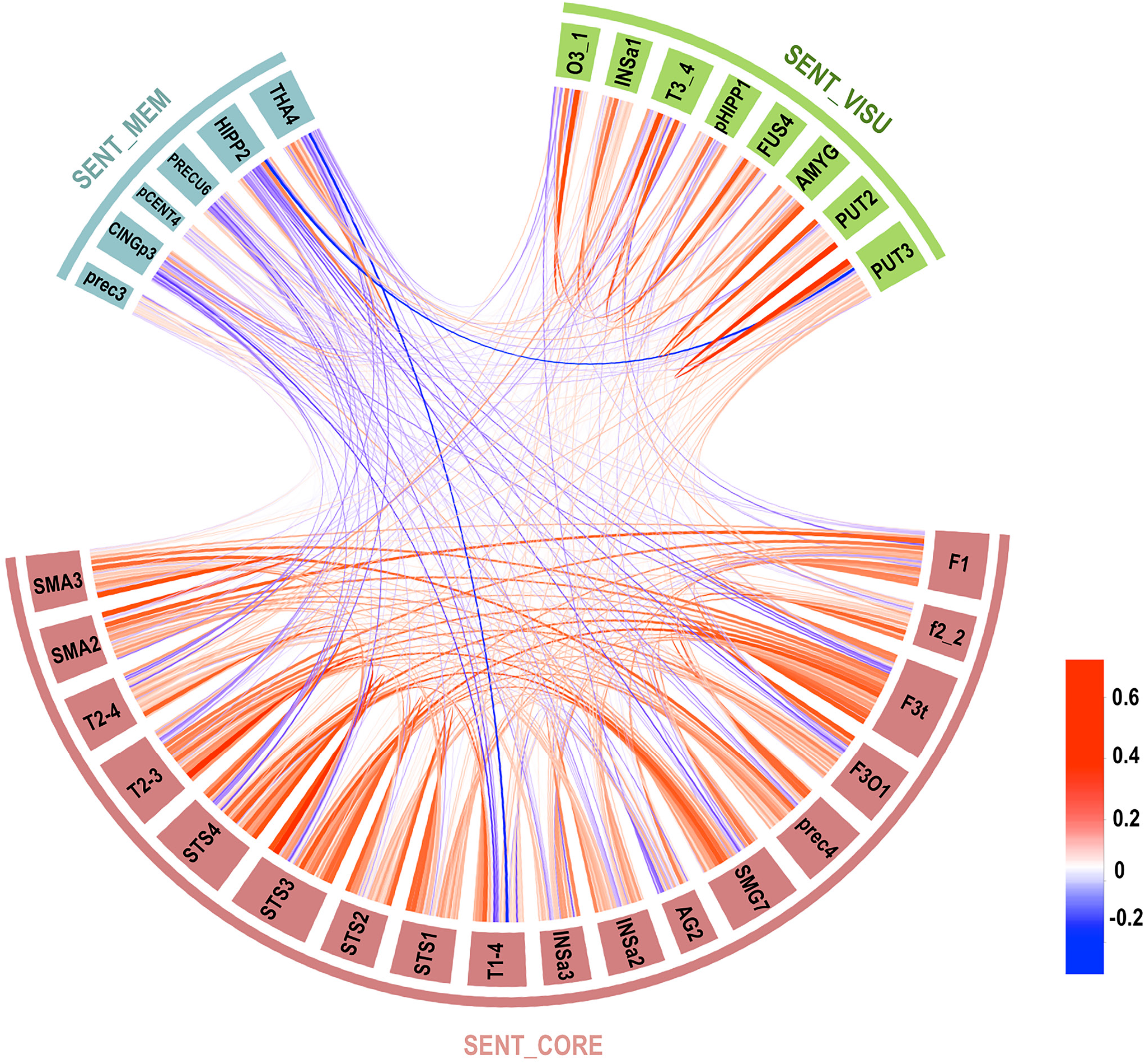
Chord diagram of the temporal correlation across each hROI composing the 3 networks averaged in the whole group. Abbreviations for hROIs of the AICHA atlas can be found in Table 2 (colour scale goes from red for positive correlation to blue for negative correlations, and the line width indicates the strength of the correlation).

## PART 3. Graph Theory Analysis of the Sent_Core Network

We applied graph theory analysis to the SENT_CORE hROI pairwise correlation matrix, including only positive correlations since the inclusion of negative correlations in graph theory analysis remains controversial (Rubinov and Sporns, 2010). Note that the graph theory analysis of intra-network communication was completed for only SENT_CORE, as the other 2 networks had too few nodes.

## Statistical analysis

### Identification of hubs using graph analysis metrics of the networks

#### Measurements of weighted centrality

We measured the degree centrality (DC) of the hROIs composing the SENT_CORE network, corresponding to the sum of the strength of the positive correlation of each node (hROI). DC can thus be interpreted as the amount of information that a given hROI receives from the hROIs to which it is directly connected; i.e., the DC measures the importance of a given hROI within its network according to the number and strength of interactions it undergoes with the other hROIs.

The betweenness centrality (BC) was also measured for SENT_CORE as defined by Opsahl et al. (Opsahl et al.). The BC of an hROI can be interpreted as the participation rate of that hROI in the set of shortest paths between any pair of nodes within the network; i.e., BC measures the dependence of the network on a specific hROI for its communication.

#### Hub definition and clustering

To discriminate hubs among SENT_CORE, we applied a combination of Sporns (Sporns et al., 2007) and van den Heuvel (van den Heuvel et al., 2010) definitions. We considered that an hROI had the properties of a hub when its DC and BC values were larger than the means plus one standard deviation of the DC and BC values of the hROI set in the network.

To assess whether the hubs identified in SENT_ CORE participated in communication with the other 2 networks or whether its communication was only intra-SENT_CORE, we calculated the participation index (pIndex) criteria as defined by Guimera (Guimera and Amaral, 2005). hROIs having the 15% highest pIndex values were considered connector hubs (i.e., between networks) (van den Heuvel and Sporns, 2011), while the other hROIs corresponded to provincial hubs, i.e., an hROI communicating with only its own network.

### Investigation of the relationship between intrinsic connectivity and activation measured during the language tasks

#### Relationships between DC and BOLD variation at the hROI level

We investigated whether an hROI exhibiting high intrinsic connectivity with other areas of the SENT_CORE was more activated during language tasks. For this, we performed a MANCOVA with repeated measures with a TASK main effect, a DC main effect and a DC by TASK interaction. Correlations values and corresponding p-values between the DC and activation were computed for each hROI and each task.

To test the specificity of this relationship, we completed similar MANCOVA with DC measurements obtained for the 185 hROIs of the AICHA atlas covering the entire left hemisphere. Therefore, the DC values were computed considering the connections of hROIs belonging to SENT_CORE with all other hROIs of the left hemisphere. This analysis made it possible to more deeply characterize whether the relationship between the DC and activation during the language tasks was specific to the essential language network intrinsic connectivity or whether this relationship was held at the hemispheric level.

#### Relationships between the DC measured in the SENT_ CORE network and BOLD variation upon pooling all hROIs and participants

To test whether the relationship previously identified between DC and BOLD signal variation for each hROI of SENT_CORE was a general property that could be extrapolated to any hROI of any participant, we applied the method proposed by Buckner for evaluating the relationship between DC and beta-amyloid accumulation in Alzheimer’s disease (Buckner et al., 2009). Correlation coefficients obtained for the 3 tasks between DC values and BOLD variations were compared using the R package “cocor” function (Diedenhofen and Musch, 2015) to determine whether there was any difference across language tasks.

## Results

### Graph analysis of SENT_CORE

#### Sample distributions of DC and BC values

DC variation across the hROIs spanned from 2.24 to 6.48 (Table 4), and the DC standard deviation was very consistent across hROIs ranging from 1.17 to 1.79.

**Table 4.** Betweenness and degree centrality of SENT_CORE hROIs. The means and standard deviations (SD) of the betweenness centrality (BC) and degree centrality (DC) were computed by averaging the BC and DC values of each participant for each SENT_ CORE hROI. For BC, the percentage of null values is based on the number of BC values at zero among the 138 subjects for one hROI. For DC, the skewness, kurtosis and Shapiro-Wilk normality test (p norm) correspond to information regarding the normality of the DC distribution for each hROI. A value above 0.05 for the Shapiro-Wilk normality test indicates that the DC was normally distributed.

|  | AICHA<br>hROI | Betweenness Centrality |  |  | Degree Centrality |  |  |  |  |
| --- | --- | --- | --- | --- | --- | --- | --- | --- | --- |
|  |  | Mean | SD | % null<br>values | Mean | SD | Skewness | Kurtosis | p norm |
| Frontal &<br>insula | prec4 | 6.88 | 7.00 | 11 | 4.94 | 1.50 | -0.07 | -0.24 | 0.87 |
|  | F1_2 | 6.30 | 5.42 | 17 | 4.64 | 1.40 | 0.28 | -0.32 | 0.27 |
|  | f2_2 | 3.33 | 4.45 | 37 | 3.35 | 1.53 | 0.60 | -0.42 | 0.0002 |
|  | F3t | 18.98 | 9.77 | 1 | 6.48 | 1.26 | -0.06 | -0.29 | 0.78 |
|  | F3O1 | 2.18 | 4.04 | 54 | 4.31 | 1.50 | 0.50 | -0.02 | 0.018 |
|  | INSa2 | 2.49 | 3.66 | 43 | 3.84 | 1.46 | 0.27 | -0.23 | 0.14 |
|  | INSa3 | 2.54 | 3.98 | 40 | 3.01 | 1.29 | 0.76 | 0.63 | 0.0005 |
| Temporal &<br>parietal | T1_4 | 0.75 | 2.19 | 77 | 3.12 | 1.47 | 0.56 | -0.20 | 0.0012 |
|  | T2_3 | 5.28 | 5.96 | 29 | 5.44 | 1.43 | 0.05 | -0.17 | 0.71 |
|  | T2_4 | 2.21 | 4.28 | 58 | 3.46 | 1.79 | 0.24 | -0.60 | 0.074 |
|  | STS1 | 2.46 | 3.57 | 41 | 4.55 | 1.52 | -0.10 | -0.18 | 0.44 |
|  | STS2 | 3.07 | 4.85 | 41 | 4.30 | 1.44 | 0.14 | -0.58 | 0.11 |
|  | STS3 | 13.04 | 9.66 | 6 | 6.32 | 1.36 | -0.36 | 0.23 | 0.088 |
|  | STS4 | 13.33 | 8.25 | 2 | 5.58 | 1.51 | 0.09 | -0.22 | 0.95 |
|  | SMG7 | 6.23 | 6.86 | 22 | 5.28 | 1.60 | -0.21 | -0.64 | 0.037 |
|  | AG2 | 0.72 | 1.65 | 71 | 2.24 | 1.17 | 0.88 | 1.11 | 0.0002 |
| Internal<br>surface | SMA2 | 2.21 | 3.58 | 46 | 4.35 | 1.33 | 0.33 | -0.09 | 0.32 |
|  | SMA3 | 2.24 | 3.60 | 49 | 4.22 | 1.28 | 0.05 | -0.36 | 0.45 |

By contrast, BC variation across the hROIs spanned from 0.72 to 18.98 (Table 4). Notably, only 3 hROIs had low numbers of BC null values across the sample of 138 participants: F3t, STS3 and STS4 (1%, 6%, and 2% null values, respectively, Table 4).

#### Hub identification and characterization

Three hROIs corresponded to the hub definition, i.e. BC and DC values above the chosen significance thresholds (mean + SD) of 10.26 and 5.55, respectively (Table 4). The first hROI (F3t) was located in the frontal lobe, and the other two were located in the posterior third of the STS (STS3 and STS4). The BC values of these 3 hubs were over 11, and their DC values were over 5.5, with F3t having the strongest values (Figure 4, Table 4). Note that no other hROI exhibited a supra-threshold value for any of the centrality indices.

**Fig. 4.**
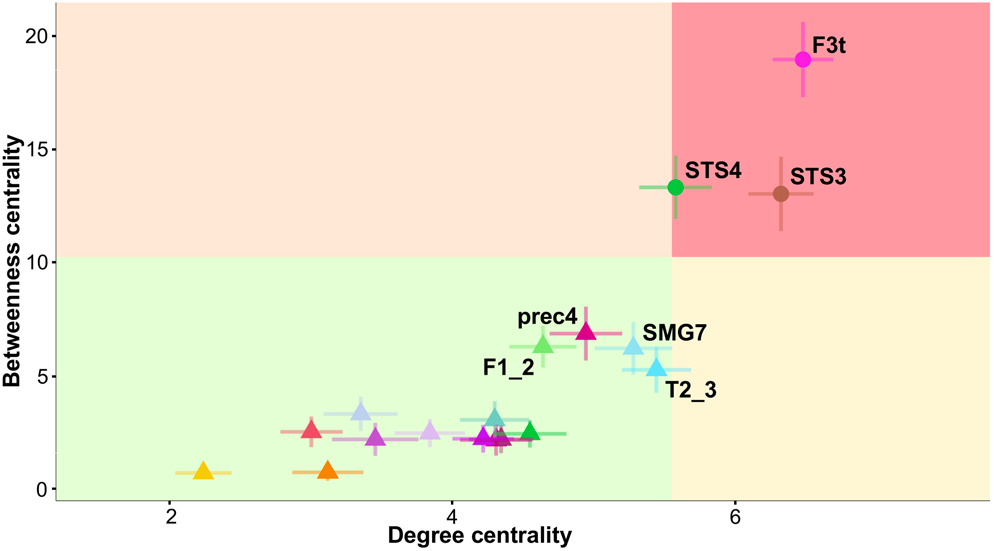
Plot of degree centrality (DC) versus betweenness centrality (BC) in SENT_CORE. The mean plus standard deviation values of DC and BC define the quadrants. hROIs located in the superior right quadrant are hubs. Abbreviations for the hROIs of the AICHA atlas can be found in Table 2.

Concerning the pIndex, hubs were defined as the top 15% of the highest index (pIndex ranging from 0.587 to 0.989). Five hROIs were thus defined as connector hubs :T2_3 (pIndex = 0.989), F3t (pIndex = 0.987), F1_2 (pIndex = 0.984), STS3 (pIndex = 0.983) and SMA2 (pIndex = 0.983).

Note that the centrality hubs F3t and STS3 were also connector hubs, meaning that they are important for both communication among the 3 different networks and for communication within the SENT_CORE network.

Note that T2_3 was a connector hub characterized by high DC and BC values (Table 4), although it did not meet the criteria to be labelled as a centrality hub (Figure 4).

### Relationship between the DC at rest and activations during the language tasks in the SENT_CORE network

#### Relationship at the individual hROI level

Using DC values computed from only SENT_CORE hROIs, we observed significant positive correlations between activations during the 3 language tasks and these DC values in 12 hROIs among the 18 constituting SENT_CORE together with a trend for PREC4 and f2_2 (Table 5). Among these hROIs, DC values were positively correlated with activations during the language tasks and the R value varied between 0.17 and 0.33. Moreover, in 8 of these 12 hROIs, there was no DC by Task interaction, meaning that the correlation between the DC and activation did not differ between the tasks. In the f2_2, INSa2, INSa3, T1_4 and SMG7 hROIs, a significant DC by Task interaction was observed. In f2_2, the interaction was due to non-significant correlation for the production task contrast, while the correlation was strong and significant for the reading and listening task contrasts. In INSa2, INSa3 and T1_4, the interaction was due to non-significant correlation for the listening task, while there were strong and significant correlations for production and reading. In SMG7, the interactions were due to a lower correlation during listening (Table 5).

**Table 5.**
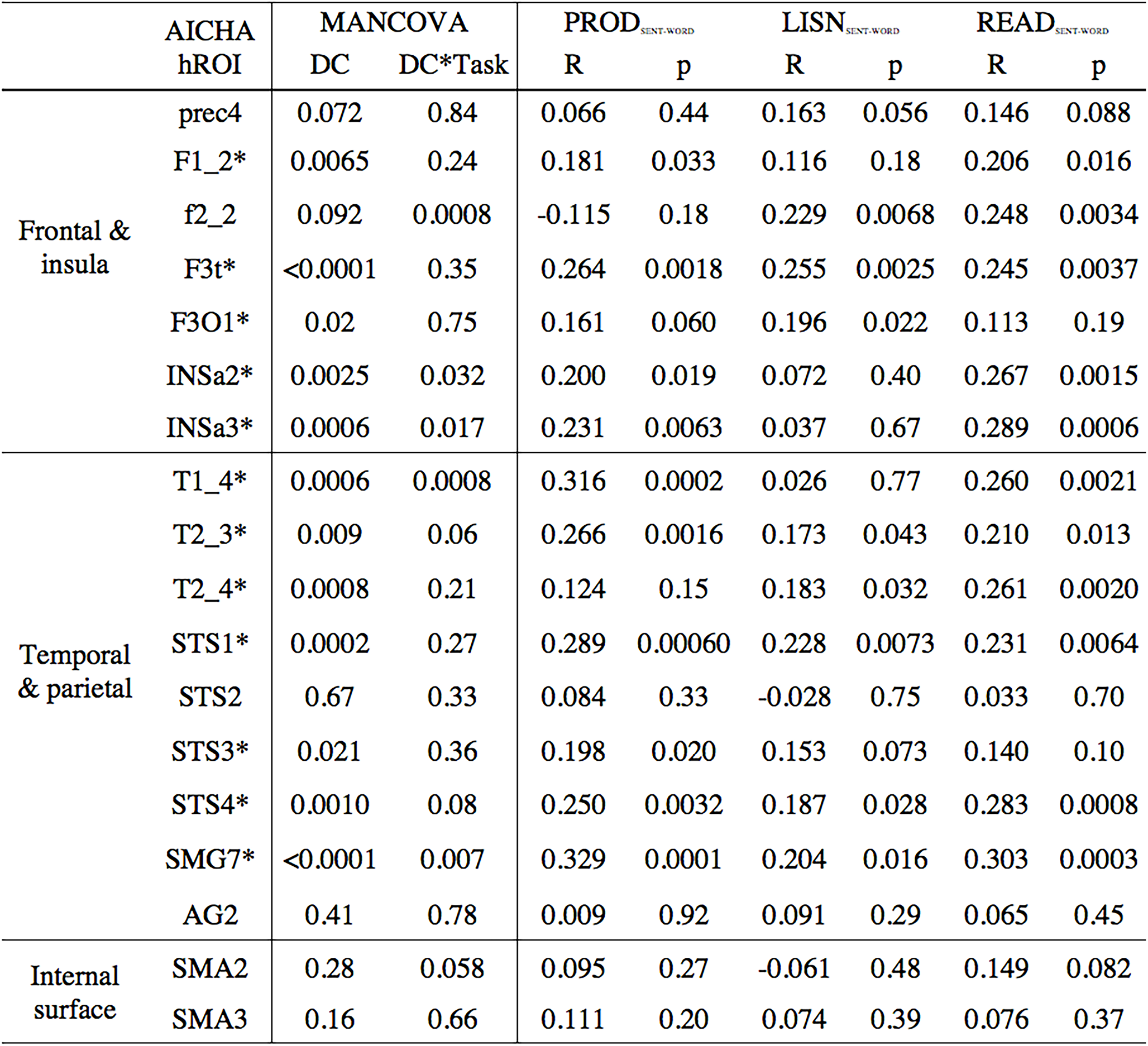
Correlation analysis between the degree centrality measured in the SENT_CORE network and the mean activation in each of the 3 language tasks. Correlations (R) were calculated within each hROI of the left hemisphere constituting the SENT_CORE network, and the DC values were calculated in the SENT_CORE network. hROIs with a star (*) are those with significant correlations between activation and DC values (p < 0.05).

The results obtained using DC values computed from the entire set of 185 left hemisphere hROIs were strikingly different. There was a significant main effect of the DC in only 2 hROIs (F3t and SMG7, see supplementary Table 2), meaning that, except for these two regions, the strength of the correlation when the DC was calculated across the hROIs of the entire hemisphere did not explain the activation variations in SENT_CORE hROIs.

#### Relationship at the global level using all participants and hROIs

There was a significant correlation between the DC values and BOLD variations measured in each of the 3 tasks when considering the 18 SENT_CORE hROIs and the 138 participants in a single analysis (Figure 5) for each task. The coefficient correlation values were 0.158, 0.216 and 0.294 for sentence production, sentence listening and sentence reading, respectively. The correlation for reading was significantly larger than that for both listening

**Fig. 5.**
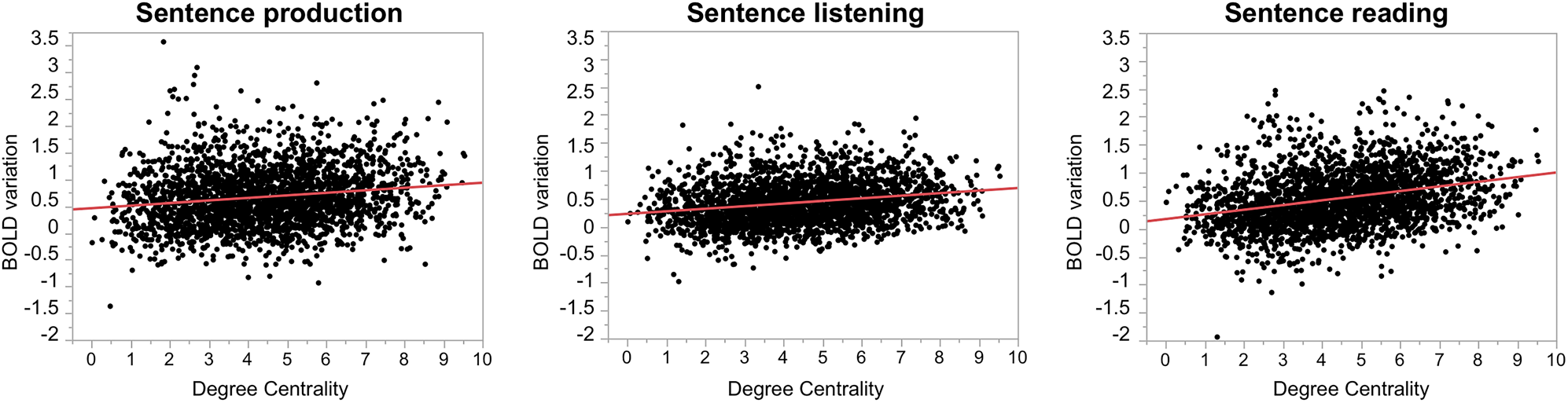
Correlation between DC values and activations in SENT_CORE across participants and across the 18 hROIs during each of the 3 language tasks. Plots of DC values and BOLD variations of the sentence minus word contrasts calculated for sentence production (left), sentence listening (middle), sentence reading (right) and degree centrality. The positive correlation coefficients (N = 138*18 = 2484) are 0.158 for sentence production, 0.216 for sentence listening, and 0.295 for sentence reading. (p = 0.0025) and production (p = 0.0075), and the latter two were not significantly different (p = 0.80).

## Summary of Results

Conjunction analysis of left activated and leftward asymmetrical hROIs in 144 right-handed participants performing three language tasks (PROD_SENT-WORD_, READ_SENT-WORD_ and LISN_SENT-WORD_) uncovered a set of 32 supramodal regions involved in lexico-syntactic processing. The hierarchical bottom-up clustering of the intrinsic connectivity between these 32 hROIs led to the identification of 3 networks, including a network of essential language areas (SENT_CORE) with strong positive correlations at rest across its 18 hROIs in more than 90% of the participants. The two other identified networks had lower inter-individual consistency, one including visual language areas at the interface between visual and syntactic processing (SENT_VISU) and the other including posterior DMN areas including posterior hippocampus (SENT_MEM). Intrinsic connectivity analysis showed that SENT_CORE was negatively correlated with SENT_MEM but was not correlated with SENT_VISU. Graph analysis metrics obtained for the SENT_CORE network revealed that F3t, STS3, and STS4 were hubs of both degree and betweenness centrality, and F3t and ST3 were also hubs of participation, meaning that these are key areas for both intra-network communication and inter-network communication between SENT_CORE and the other 2 networks. Importantly, a positive correlation across individuals was observed between the DC measured at rest and the strength of activation in most SENT_CORE regions, meaning that participants with higher DC values in a given region had higher activations than participants with lower DC values. Moreover, such a positive correlation between the DC and activation was still significant when all regions of all participants in the 3 tasks were pooled, meaning that this was true regardless of the cortical area considered.

## Discussion

### Methodological issue

In this study, we selected right-handers from the BIL&-GIN database because we previously demonstrated that these participants have a left hemisphere dominance for language at both the group level (Tzourio-Mazoyer et al., 2016) and the individual level (Zago et al., 2017), with only 5 (3%) of the 144 participants having a co-dominant right hemisphere. This sample group is optimal for selecting areas specific for sentence processing based on a conjunction of activations and leftward asymmetries. In addition, the inclusion of a fairly considerable number of participants (N = 144) provided us a high sensitivity for detecting supramodal sentence areas while minimizing the risk of overlooking some.

However, we must underline that the present atlas is not all-inclusive. First, we selected map regions involved in only high-order language processing and lexi-co-syntactic processing. Using the list of familiar words as the reference condition, we removed the dorsal route of language, including the phonological loop, responsible for articulation and sound to articulation mapping (Saur et al., 2008, Rauschecker and Scott, 2009) In addition, the regions selected herein focused on the left hemisphere and did not account for right hemisphere-specialized aspects of sentence processing, such as emotional prosody (Beaucousin et al., 2007, Hurschler et al., 2012) and context processing (Grindrod and Baum, 2003, Ferstl et al., 2005). Second, the presence of susceptibility artefacts combined with averaging the large number of participants led to incomplete mapping of the inferior part of the temporal lobe, prohibiting us from documenting some areas, such as the basal language area in the anterior part of the fusiform gyrus. This essential language area, first identified using deep electrical recordings (Nobre et al., 1994), has been shown with positron emission tomography (PET) to be activated during both the production and auditory comprehension of language (Papathanassiou et al., 2000).

Third, small size regions may also be lacking in this atlas since we provided data at the hROI scale rather than at the voxel scale.

Concerning the clustering methods of correlation values at rest, a perfect match was observed between the Ward and average clustering methods (see supplementary material Table 1), and a good score was obtained with the Gaussian Mixture Model for global clustering at the 32 hROI levels. The weakest score was that of the k-means method, and such a difference in clustering observed with k-means compared to that in the other 3 methods is consistent with the fact that, as reported by Thirion et al., k-means forms clusters spatially close and connected but with poor reproducibility using the sample studied. By contrast, hierarchical clustering using Ward’s method, which we selected to segregate the networks, was reported to create connected clusters that are highly reproducible using the studied samples (Thirion et al., 2014).

### A large set of supramodal language areas is involved in sentence processing tasks

We carefully designed each of the language tasks such that joint analyses were possible; the design was identical in the 3 tasks, and we chose to make them close enough to allow comparisons and conjunctions in terms of the number of words or the complexity of sentences. As mentioned above, the use of a high-level verbal reference task for controlling the involvement of primary areas (auditory, visual and motor) and removing phonological and automatic word processing kept the lexico-syntactic aspects common to all three tasks.

The first set of 32 hROIs provides left hemispheric regions that are dedicated to the monitoring and completion of tasks based on sentence processing. Although not all regions can be considered essential language areas, all were determined to be modulated by the verbal material with which they are associated (left activation and leftward asymmetry) and are thus part of an extended language network functioning during language tasks.

Unsupervised hierarchical clustering based on the resting-state connectivity between hROIs successfully segregated 3 different networks, including networks hosting core language, visual areas, and posterior areas of the DMN and posterior hippocampus. Within the systems to which they belonged, these networks hosted areas dedicated to the interaction/interface with language systems. For example, the current analysis extracted from among the visual areas involved in picture processing those areas specifically dealing with picture-sentence meaning integration.

### Sentence comprehension essential network (SENT_CORE)

Clustering the resting-state correlation between these 32 hROIs allowed the discrimination of SENT_CORE, a network of 18 strongly and positively correlated hROIs, including frontal and temporo-parietal hROIs located on the lateral surface of the left hemisphere and anterior insula areas. In particular, SENT_CORE included areas of the antero-posterior language networks, named in reference to the Broca-Wernicke model in aphasia literature and reported with consistency in meta-analyses of healthy individuals mapped during language tasks (see Figure 6, (Vigneau et al., 2006, Price, 2010, 2012). Note that SENT_CORE was the largest network in terms of volume (in mm3), as it included more than half of the hROIs (18/32), all of which were strongly activated and leftward asymmetrical.

**Fig. 6.**
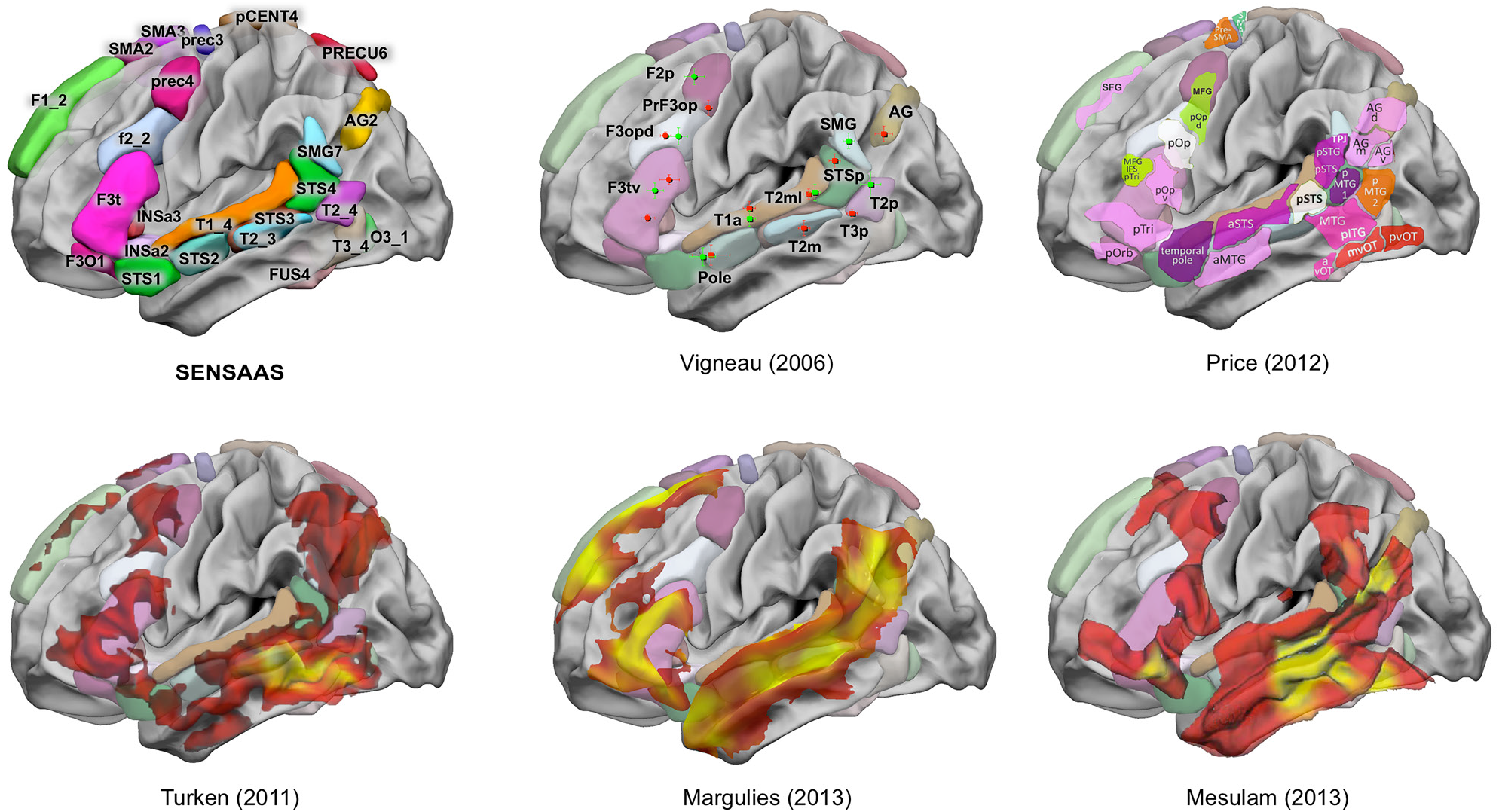
Schematic comparison of SENSAAS with the literature. This figure shows the results of neuroimaging meta-analyses and clinical studies superimposed on the hROI of SENSAAS of the lateral surface of the left hemisphere of the BIL&GIN display template. In the first row: left SENSAAs hROIs of the left hemisphere lateral surface; middle: clusters of the meta-analysis of semantics (red) and sentence processing (green) adapted from Vigneau (2006) with their labels: right: schematic representation of the meta-analysis of language-related activation studies (adapted from Price (2012); sentence: purple; semantics: light and dark pink; visual: red; word retrieval: green; integration: white). In the second row, left: functional connectivity of the left middle temporal gyrus centred on the site where lesion results in deep aphasia (orange, adapted from Turken 2011); middle: functional connectivity from a seed centred on the left inferior frontal gyrus (BA 45, red, adapted from Margulies 2013); right: zones of atrophy observed when pooling all types of PPA (orange, adapted from Mesulam 2013).

In the following, we discuss the potential roles of the identified areas in relation to the literature. However, it is now acknowledged that, apart from very specific regions where a lesion can be closely associated with a specific defect, the role of a given area documented with functional imaging must be understood as the combination of its functional properties with those of the regions with which it constitutes a network to complete a given cognitive task. For example, prec4 is not part of the regions commonly labelled as “language areas”. The present work shows that prec4, located at the junction between the precentral and middle frontal gyrus, is both strongly activated and leftward asymmetrical in the 3 sentence tasks. Language meta-analyses have reported prec4 as part of the language areas involved in lexico-syntactic processing (named F2p in Vigneau et al., 2006, Figure 6), and in word selection and hierarchical sequencing (named dPrec in Price, 2010, Figure 6). Applying Neurosynth to prec4 coordinates (× = −42.2, y = 0.7, z = 50, Table 3) with an association test reveals the greatest number of studies with the terms «sentence», «comprehension», «language», and «sentences», followed by «eye» and «premotor». Jouen et al. (Jouen et al., 2018) propose that prec4 is “involved in the understanding of actions during verbal and non-verbal tasks”. In the present protocol, the sentences involved human actions, closely consistent with that role proposed by Jouen et al. (2015). Using a network approach, Saur et al. (Saur et al., 2008) found that prec4 belongs to the sentence comprehension functional network. In Glasser’s atlas, prec4 corresponds to “language area 55b” (Glasser et al., 2016) and overlaps the posterior part of the “rostro-ventral module” described by Genon et al. (Genon et al., 2018). In this last work based on peaks meta-analysis, this module is connected with the inferior frontal gyrus, orbital frontal and inferior parietal as prec4 in the present work. The present study further demonstrates that prec4 is involved in a supramodal manner during sentence processing, is strongly leftward asymmetrical, and is strongly and positively connected at rest with the network of areas we named SENT_CORE that hosts essential language areas. These findings are consistent with the fact that prec4 is part of the areas that are conjointly atrophied in patients suffering from nonfluent primary progressive aphasia (PPA) (Mesulam et al., 2014) Figure 6).

Indeed, in the frontal lobe, SENT_CORE includes the whole inferior frontal gyrus (F3t and F3O) corresponding to Broca’s area according to most authors, whose lesion is responsible for conversational deficits, as is also the case for the anterior insula (INSa2, INSa3) (Borovsky et al., 2007). The posterior part of the inferior frontal sulcus (f2_2), also part of SENT_CORE, has been underlined as an area involved in lexico-syntactic processing (named F3opd in Vigneau et al., 2006, Figure 6). Price also targeted f2_2 as being involved in word selection and hierarchical sequencing (named mFG in Price, 2010, Figure 6). Another indication of the important role of f2_2 in language is that together with F3t and prec4, it is the location of atrophy in non-fluent PPA ((Mesulam et al., 2014) see figure 6).

In the medial part of the frontal lobe, SENT_CORE includes both preSMA and the superior frontal gyrus (here, SMA2, SMA3 and F1_2), which have been reported in tasks involving sentences dealing with characters (Herve et al., 2012), as in the present paradigm. The activation of these areas has been attributed to processing the social aspects of verbal material (Ferstl and von Cramon, 2002). As discussed below, these medial frontal areas are strongly connected with the posterior part of the middle temporal gyrus (Turken and Dronkers, 2011). Notably, these medial frontal areas are also the sites of atrophy in PPA (Wilson et al., 2010, Tetzloff et al., 2017).

In the temporal lobe, SENT_CORE included the STS together with the posterior part of the middle and superior temporal gyri, extending to the angular and supramarginal gyri. This characterization is consistent with the proposal by Price that these areas are involved in amodal semantic combinations, a process common to the 3 sentence tasks (Price, 2010). A recent work on the time course of sentence processing areas has also shown that the posterior temporal lobe, here corresponding to STS3 and T2_3, is involved in lexico-syntactic processing, with the processing of individual words in relation to the syntactic structure (Matchin et al., 2018). These posterior temporal areas have been documented to be essential language areas since the lesion of each results in specific deficits in sentence comprehension (Dronkers et al., 2004). In particular, lesion of the posterior parts of the middle temporal gyrus results in deep aphasia due to a loss of word meaning (Dronkers et al., 2004), consistent with the specific atrophy observed in logopenic PPA (Wilson et al., 2010). STS4, located dorsally to STS3 and T2_3, is involved in sentence-level high-order processing, and Matchin, in his work on the time course of sentence processing investigated with MEG, proposed that “increased activation at the end of the sentence suggests a response associated with the interpretation of the sentence” (Matchin et al., 2018). The integration gradient of sentence meaning from lexical to conceptual progresses posteriorly with SMG7, which corresponds to the area labelled STSp in Vigneau et al.’s meta-analysis (Figure 6). Vigneau et al. reported the following: “STSp role seems to process the semantic integration of complex linguistic material. This statement comes from the observation that it is recruited when subjects listen to coherent rather than syntactically or pragmatically incoherent sentences (Kuperberg et al., 2000, Luke et al., 2002), and it is involved in context processing and syntactic generation—more activated when subjects have to choose between two words to end a sentence or have to generate the final word of a sentence (Kircher et al., 2001). STSp activity is very likely related to the linkage of linguistic structure to meaning: it is more activated when sentences are linked as dialogue (Homae et al., 2002) or syllogisms (Goel et al., 1998) than when they are unlinked. It is also more activated during text comprehension, either presented auditory (compared to reverse speech (Kansaku et al., 2000, Crinion et al., 2003) or words (Jobard et al., 2007)) or visually (compared to words (Jobard et al., 2007) or pseudo-word reading (Vingerhoets et al., 2003))”. Price made an alternative proposal in her meta-analysis in 2000 stating that the equivalent of SMG7 (vSMG and pPT) is activated by sentence processing, particularly when the sentences increase in difficulty, and she therefore suggested that SMG7 is involved in subvocal articulation. The present results demonstrating that SMG7 is involved and leftward asymmetrical not only during production and reading but also during sentence listening, which was cited as a easy task by the participants, suggests its involvement in meaning elaboration. The angular gyrus is the final location where this integration towards conceptual knowledge operates dorsally. Known to be involved in lexico-semantic processing, as identified by language task-induced activation studies ((Vigneau et al., 2006, Price, 2010), see Figure 6), the angular gyrus is specifically involved in conceptual knowledge as shown by neuropsychological studies. Lesion of this area leads to the inability to associate a sound or image related to the same concept (Saygin et al., 2004), and inactivation of the angular gyrus in the left hemisphere induces a deficit in semantic integration (Price et al., 2016). The angular hROI identified herein corresponds to Wang’s C5 and C6 parcels, involved in both language and theory-of-mind tasks (Wang et al., 2017). This last observation confirms that the role of the AG includes the interfacing between sentence processing and the understanding of human actions, a process that is part of the present sentence tasks.

The similitude between the areas composing the SENT_CORE network and the regions showing atrophy in all types of PPA ((Mesulam et al., 2014), see Figure 6) is striking and a key element evidencing that SENT_CORE contains essential language areas, although the anterior part of the inferior temporal gyrus hosting the basal temporal language area (Nobre et al., 1994, Papathanassiou et al., 2000) is lacking for the methodological reasons discussed above.

Finally, the language functional network, which shares a common molecular basis of interaction highlighted by Zilles et al. using neurotransmitter receptor fingerprints (Zilles et al., 2015), overlaps the hROIs of SENT_CORE, apart from AG2. Based on Zilles et al. report, we can hypothesize that the two other sentence-related networks we have identified are likely to have different fingerprints from those of SENT_CORE. However, because the hROIs of these networks were selected as both activated and leftward asymmetrical in the three sentence tasks, it would be of interest to investigate what subset of fingerprints the hROIs of SENT_VISU and SENT_MEM may share in common with SENT_CORE to deepen our understanding of network interactions.

### Hubs of SENT_CORE

The computation of betweenness and degree centralities allowed us to identify 3 hubs within SENT_CORE: one in the inferior frontal gyrus (F3t) and two others along the posterior part of the temporal cortex (STS3 and STS4). These high centrality figures demonstrate that these hROIs play a central role in communications with other parts of SENT_CORE, and the participation indices of F3t and STS3 further indicate their key role in communication with the other networks identified. Such properties are consistent with the definition of epicentres proposed by Mesulam (Mesulam et al., 2014), and these regions can be considered essential for sentence/test comprehension independently of the modality. According to Mesulam, the network epicentre specializes in a specific behavioural component, which is language in this study, and the destruction of transmodal epicentres causes global impairments. In fact, as noted above, these hROIs overlap the middle temporal gyrus targeted in the aphasics investigation by Dronkers (Dronkers et al., 2004) as well as the regions showing the highest atrophy in all types of PPA (Mesulam et al., 2014), confirming that they do correspond to epicentres. In addition, these hubs are distributed in the anterior and posterior cortices, constituting an antero-posterior loop across areas belonging to the same hierarchical level in terms of cortical organization, consistent with Fuster’s model for cognition (Fuster and Bressler, 2012).

Other studies involving functional connectivity at rest have shown the strong connection of the left inferior frontal gyrus at rest with all SENT_CORE areas ((Margulies and Petrides, 2013), Figure 6). The similarity between the regions found by these authors when seeding the posterior part of the middle temporal gyrus (here, STS3 and STS4) and the areas constituting SENT_CORE is also clear ((Turken and Dronkers, 2011), Figure 6). These two observations are consistent with the present observation that these regions are hubs for sentence processing and, more generally, the language comprehension network. Such a framework is also thought to correspond to the Broca-Wernicke language model, with F3t serving as Broca’s area and STS3 and STS4 serving as regions involved in the supramodal integration of meaning, consistent with the location of posterior areas leading to comprehension deficits (Pillay et al., 2017). These posterior STS areas are also considered by Binder as areas supporting meaning integration during sentence comprehension (Binder, 2017). Considering that a left deficit in this area leads to deficits in language comprehension, we propose to label it Wernicke’s area, although as reviewed by Binder, such a definition is different from the location currently proposed for Wernicke’s area in the superior temporal gyrus. In the present work, closely adhering to an anatomo-functional definition in reference to deficits in comprehension associated with Wernicke’s aphasia (Dronkers et al., 2004, Binder, 2015, 2017), we propose that F3t, STS3 and STS4, regions strongly activated and asymmetrical during sentence processing in different modalities and hubs of the SENT_CORE network, are the epicentres of sentence comprehension.

### Other areas contributing to sentence tasks

Because there was a low inter-individual consistency in the segregation of the networks labelled SENT_VISU and SENT_MEM, we will discuss the potential role of the 14 hROIs that were not part of SENT_CORE together.

Because each event of the sentence tasks included task monitoring, such as shifting between the word list and sentence tasks when a picture was presented or providing a motor response at the end of sentence processing, the involvement of executive areas was expected. Conjunction analyses indeed revealed that, in addition to the anterior insula at play in the 3 tasks (see above), putamen hROIs were involved and are a key neural support of executive functions and task monitoring (Monchi et al., 2006, Sefcsik et al., 2009). Moreover, a recent meta-analysis involving connectivity modelling showed that the left and right putamen areas are different in terms of their respective co-activations, which are specifically co-involved in language areas (Viñas-Guasch and Wu, 2017).

The posterior cingulate, precuneus, and paracentral lobule together with the posterior hippocampus are part of the DMN, which has been shown to be involved in both episodic thinking and processing of “self”. At rest, these areas constituted a network that had a very strong and negative correlation with SENT_CORE, confirming that, although they belong to different networks, they are related at rest. Task-induced activation studies have demonstrated that posterior DMN regions are part of mind-reading areas together with the angular gyrus (Spreng et al., 2009). Activated and leftward lateralized during language tasks, this set of areas likely interacts with SENT_CORE during language tasks to process the sentence content dealing with social interactions (Herve et al., 2012). The angular gyrus is a likely candidate for such an interaction since it segregated with SENT_MEM areas in two clustering methods and is involved in both language and theory-of-mind tasks (Wang et al., 2017).

The fact that these posterior areas were segregated with the posterior hippocampus, involved in both the actual perception and the encoding of scenes (Zeidman and Maguire, 2016), suggests that the SENT_MEM network is also involved in the image processing of the drawings, a component common to the three sentence tasks. In fact, all participants performed well in the recall of these sentences (more than 9 out of 10 images recalled).

Finally, three hROIs were located along the occipito-temporal junction. The posterior part of the inferior temporal lobe (T3_4), the inferior occipital gyrus (O3_1) and the mid-fusiform gyrus (FUS4), with activity and leftward asymmetry,, are likely related to the processing of the sentence content in relation to the one-second drawing presentation (minus the scramble version of the word-list reference tasks) common to all tasks. In her meta-analysis, Price considered these areas to be involved in direct visual combinations (Price, 2010), consistent with the design of the present paradigm wherein the participants dealt with images related to sentence content regardless of the sentence task modality and the fact that the mean value and asymmetry of this network did not vary with the language tasks. Furthermore, in her 2012 review, Price considered these areas to be involved in sentence processing depending on the task demand (Price, 2012). FUS4 is a region of the ventral route that corresponds to the visual word form area (Mellet et al., 2018). It is notable that this region was activated and leftward asymmetrical during the 3 tasks and segregated with the other visual regions at rest. Such a behaviour is in accordance with Price and Devlin’s proposal that the mid-fusiform area is involved not only in word and visual processing but is a multimodal area whose function is determined by the set of interacting areas (Price and Devlin, 2003). FUS4 involvement in the present study is likely to be in the association between the meaning conveyed by the pictures and the sentences and appears to be at play whatever the sentences’ modality. The fact that FUS4 did segregate with visual areas rather than sentence areas at rest supports this hypothesis (Figure 2). In fact, visual hROIs were not correlated with SENT_CORE networks, meaning that they do not exchange information at rest. However, these areas involved in visual processing, as well as the other hROIs that did not constitute a network at rest with sentence core areas, showed a strong leftward activation during the language tasks, suggesting that their involvement during sentence is, at least in part, related to top-down influences from language networks.

### The degree of centrality measured in SENT_ CORE explains the activation variability during the 3 language tasks

To our knowledge, this study is the first to report a positive correlation between DC and task-induced activation values. Considering that the correlations between DC values and activation were positive for all hROIs, we suggest that the mechanism underlying this result may be that regions more highly connected within the SENT_CORE network are more highly recruited during language tasks. Such an hypothesis is supported by a very recent work conducted with electrocortical stimulation that demonstrated that critical language sites have significantly higher connectivity (Rolston and Chang, 2018). It is also important to note that for 8 of the 13 hROIs, no significant effect of the nature of the task on the correlation strength was observed: the positive correlation did not differ regardless of whether the participants produced, read or listened to sentences compared to listing words. Such a relationship was also observed when plotting all SENT_CORE regions of all subjects, as reported by others in a different context (Buckner et al., 2009), meaning that the relationship between resting-state intrinsic connectivity and activation strength remains regardless of the participant and cortical area being considered. In Buckner’s study, the authors hypothesized that beta amyloid accumulates in high DC regions because their high metabolism makes them more vulnerable to the disease, consistent with the report that the DC calculated at the voxel level in the entire hemisphere is correlated with cerebral blood flow values (Liang et al., 2013). Similarly, DC values computed at the hROI level in the SENT_CORE network may also indicate the metabolism of language areas in a given individual, which may be of interest in the evaluation of pathological states. Mesulam et al. reported the selective atrophy of right hemisphere areas in a PPA patient with right hemisphere dominance for language, an observation suggesting that language networks are specifically targeted by this illness (Mesulam et al., 2005). DC measurements in SENT_CORE may thus be a valuable index with which to evaluate inter-individual variations in language area activities in relation to anatomical and clinical patterns in such pathologies. Previous investigations dealing with the relationships between tasks and the resting state have compared the functional connectivity during cognitive tasks with that measured during the resting state (Cole et al., 2014, Gerchen and Kirsch, 2017), as reviewed by Wig (Wig, 2017). Reports more closely related to the approach used herein have compared resting-state connectivity and hemispheric activation asymmetries obtained during language production in healthy individuals (Joliot et al., 2016) and epileptic patients (Doucet et al., 2014), or during story listening (Raemaekers et al., 2018). Other investigations have found correlations between the task-induced and intrinsic connectivity asymmetries measured in selected sets of ROIs (Liu et al., 2009), such as the set of regions involved in a semantic decision task (Wang et al., 2014). Seeding specific language areas, a previous study involving epileptic patients demonstrated that the functional connectivity measured at rest was correlated with lateralization indices measured during language tasks. In this last work, the intrinsic connectivity most strongly correlated with asymmetry of task-induced fMRI was between the inferior frontal and temporo-parietal language areas, all regions constituting SENT_CORE (Teghipco et al., 2016). The set of consistent findings across the latter studies show that asymmetries of intrinsic connectivity partially explain the variability in activation asymmetries measured during language tasks. However, to our knowledge, there are no previous reports of a relationship between intrinsic DC and activation strength, probably because such a relationship is lacking when comparing whole-brain intrinsic connectivity to variations in activity triggered by cognitive tasks that are underpinned by specific networks. In fact, in the present study, this relationship was observed when the DC computation considered only the 18 hROIs of the SENT_CORE network and disappeared when the DC computation included all of the left hemisphere hROIs. This is an important observation because it underlines the necessity of exploring the properties of intrinsic connectivity within specific networks rather than at the whole-brain level.

Considering that the rs-fMRI acquisition was completed on average 11 months before the language fMRI session herein, it was surprising that the rs-fMRI-derived DC values explained up to 11% of the variance in the language fMRI-derived activation amplitudes in 12 of the 18 SENT_CORE network regions. Thus, DC values at rest in regions constituting SENT_CORE can be considered as proxies of their potential involvement during sentence processing. However, to generalize this observation, we need to both investigate how the SENT_CORE DC is modified in individuals atypical for language and to confirm that such a relationship between the DC value during the resting state and activation strength also exists for networks supporting other cognitive domains, specifically the attentional system, in the same participant.

## CONCLUSION

Based on the fMRI analysis of 3 language tasks performed by 144 healthy adult right-handers combined with the analysis of intrinsic resting-state connectivity in 138 of the participants, we propose a SENSAAS atlas of high-order sentence processing areas. This atlas includes 32 regions decomposed into 3 networks, including one (SENT_CORE) specifically composed of essential areas for sentence reading, listening and production. This atlas also contains the features of these 3 networks, a graph analysis of the intrinsic connectivity of regions that compose SENT_CORE (their degree centrality values that correlate with their strength of activation during the language tasks) as well as the relationships across the different language networks at rest. Such a positive correlation between the DC at rest and the language task-induced activation amplitude in the left hemisphere language network opens the way for investigating participants with language pathologies or population neuroimaging studies searching for the genetic basis of language by analysing only resting-state acquisition. Finally, the methodology we applied, identifying regions from activation studies for selecting the networks at play, advanced the specificity of resting-state graphical analysis and shed light on the relationships between resting-state and task-related networks.

## ACKNOWLEDGEMENTS

The authors thank Daniel Margulies for having provided images from his work to complete the figure 6 and enrich the discussion.

## ETHICS STATEMENT

### Funding

This work was supported by a grant from the FLAG-ERA Human Brain Project 2015 (ANR-15-HBPR-0001-03-MULTI-LATERAL) awarded to FC, BM, NTM, and MJ, the Ginesis-Lab project (ANR 16 LCV2 0006 01), a grant awarded to MJ, BM, NTM, EM, FC, and LP, and by a research contract from CEA to LL.

### Conflict of Interest

The authors declare that they have no conflict of interest.

### Research involving humans

### Ethical approval

The study has been approved by the Basse-Normandie research ethics committee and was performed in accordance with the ethical standards as laid out in the 1964 Declaration of Helsinki and its later amendments or comparable ethical standards.

### Informed consent

Informed written consent was obtained from all individual participants included in the study.

## SUPPLEMENTARY MATERIALS

**Supplementary Table 1.**
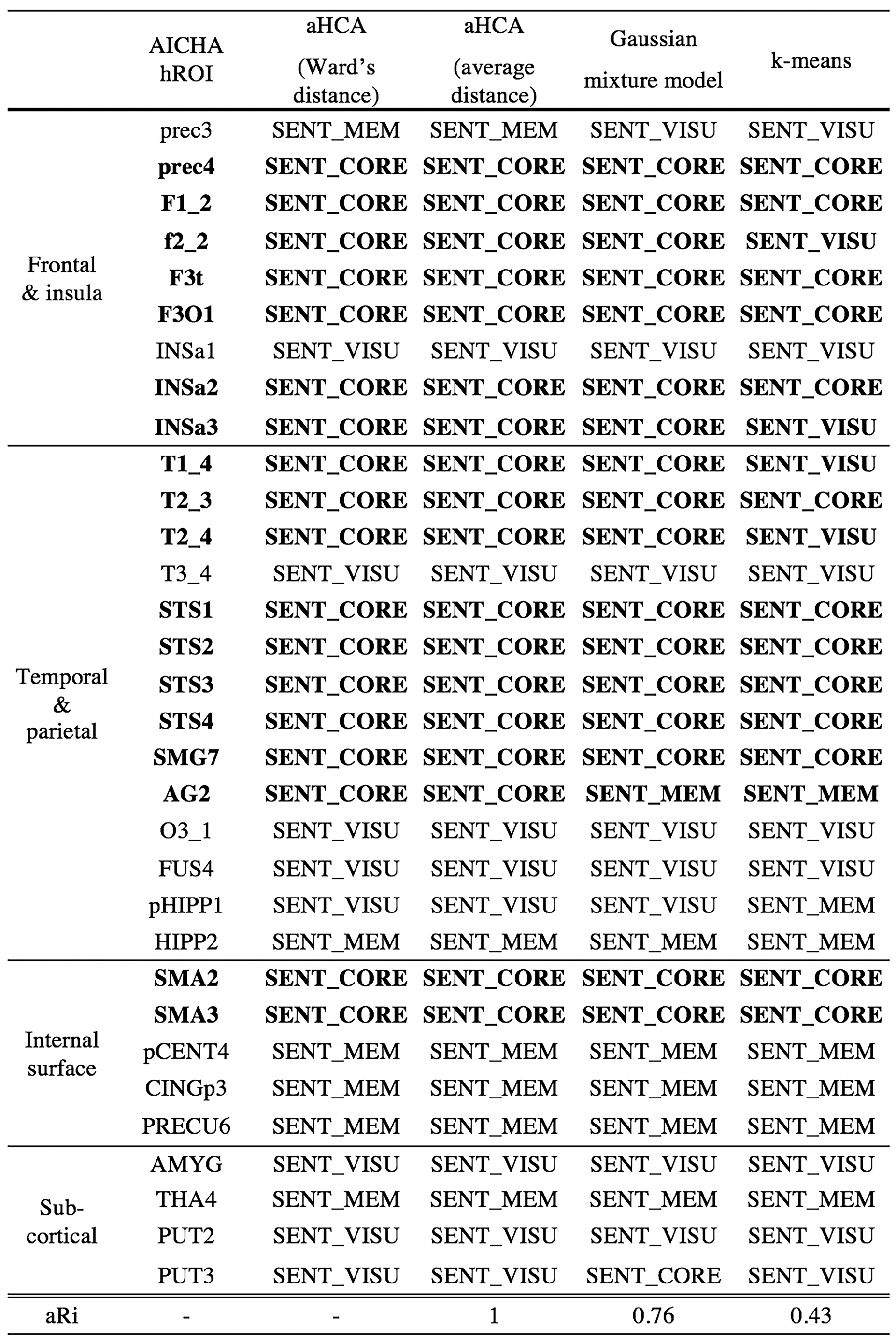
Comparison of 4 different clustering methods (agglomerative Hierarchical Cluster Analysis with Ward’s and average distance method, Gaussian mixture model and k-means) applied on the set of 32 hROIs co-activated and co-leftward asymmetrical during the sentence tasks. In aHCA, we chose the number of clusters that fulfilled a maximum of indices amongst the 30 statistical indices of the R library “NbClust”. The number of clusters in Gaussian mixture model and k-means was set according to the number of clusters find in aHCA. The last line: “aRi” (adjusted Rand index), corresponds to the classification comparison between the aHCA with Ward’s method and the 3 others.

|  | AICHA<br>hROI | aHCA<br>(Ward's<br>distance) | aHCA<br>(average<br>distance) | Gaussian<br>mixture model | k-means |
| --- | --- | --- | --- | --- | --- |
| Frontal<br>& insula | prec3 | SENT_MEM | SENT_MEM | SENT_VISU | SENT_VISU |
|  | **prec4** | **SENT_CORE** | **SENT_CORE** | **SENT_CORE** | **SENT_CORE** |
|  | **F1_2** | **SENT_CORE** | **SENT_CORE** | **SENT_CORE** | **SENT_CORE** |
|  | **f2_2** | **SENT_CORE** | **SENT_CORE** | **SENT_CORE** | **SENT_VISU** |
|  | **F3t** | **SENT_CORE** | **SENT_CORE** | **SENT_CORE** | **SENT_CORE** |
|  | **F3O1** | **SENT_CORE** | **SENT_CORE** | **SENT_CORE** | **SENT_CORE** |
|  | INSa1 | SENT_VISU | SENT_VISU | SENT_VISU | SENT_VISU |
|  | **INSa2** | **SENT_CORE** | **SENT_CORE** | **SENT_CORE** | **SENT_CORE** |
|  | **INSa3** | **SENT_CORE** | **SENT_CORE** | **SENT_CORE** | **SENT_VISU** |
| Temporal<br>&<br>parietal | **T1_4** | **SENT_CORE** | **SENT_CORE** | **SENT_CORE** | **SENT_VISU** |
|  | **T2_3** | **SENT_CORE** | **SENT_CORE** | **SENT_CORE** | **SENT_CORE** |
|  | **T2_4** | **SENT_CORE** | **SENT_CORE** | **SENT_CORE** | **SENT_VISU** |
|  | T3_4 | SENT_VISU | SENT_VISU | SENT_VISU | SENT_VISU |
|  | **STS1** | **SENT_CORE** | **SENT_CORE** | **SENT_CORE** | **SENT_CORE** |
|  | **STS2** | **SENT_CORE** | **SENT_CORE** | **SENT_CORE** | **SENT_CORE** |
|  | **STS3** | **SENT_CORE** | **SENT_CORE** | **SENT_CORE** | **SENT_CORE** |
|  | **STS4** | **SENT_CORE** | **SENT_CORE** | **SENT_CORE** | **SENT_CORE** |
|  | **SMG7** | **SENT_CORE** | **SENT_CORE** | **SENT_CORE** | **SENT_CORE** |
|  | **AG2** | **SENT_CORE** | **SENT_CORE** | **SENT_MEM** | **SENT_MEM** |
|  | O3_1 | SENT_VISU | SENT_VISU | SENT_VISU | SENT_VISU |
|  | FUS4 | SENT_VISU | SENT_VISU | SENT_VISU | SENT_VISU |
|  | pHIPP1 | SENT_VISU | SENT_VISU | SENT_VISU | SENT_MEM |
|  | HIPP2 | SENT_MEM | SENT_MEM | SENT_MEM | SENT_MEM |
| Internal<br>surface | **SMA2** | **SENT_CORE** | **SENT_CORE** | **SENT_CORE** | **SENT_CORE** |
|  | **SMA3** | **SENT_CORE** | **SENT_CORE** | **SENT_CORE** | **SENT_CORE** |
|  | pCENT4 | SENT_MEM | SENT_MEM | SENT_MEM | SENT_MEM |
|  | CINGp3 | SENT_MEM | SENT_MEM | SENT_MEM | SENT_MEM |
|  | PRECU6 | SENT_MEM | SENT_MEM | SENT_MEM | SENT_MEM |
| Sub-cortical | AMYG | SENT_VISU | SENT_VISU | SENT_VISU | SENT_VISU |
|  | THA4 | SENT_MEM | SENT_MEM | SENT_MEM | SENT_MEM |
|  | PUT2 | SENT_VISU | SENT_VISU | SENT_VISU | SENT_VISU |
|  | PUT3 | SENT_VISU | SENT_VISU | SENT_CORE | SENT_VISU |
| aRi | - | - | 1 | 0.76 | 0.43 |

**Supplementary Table 2.**
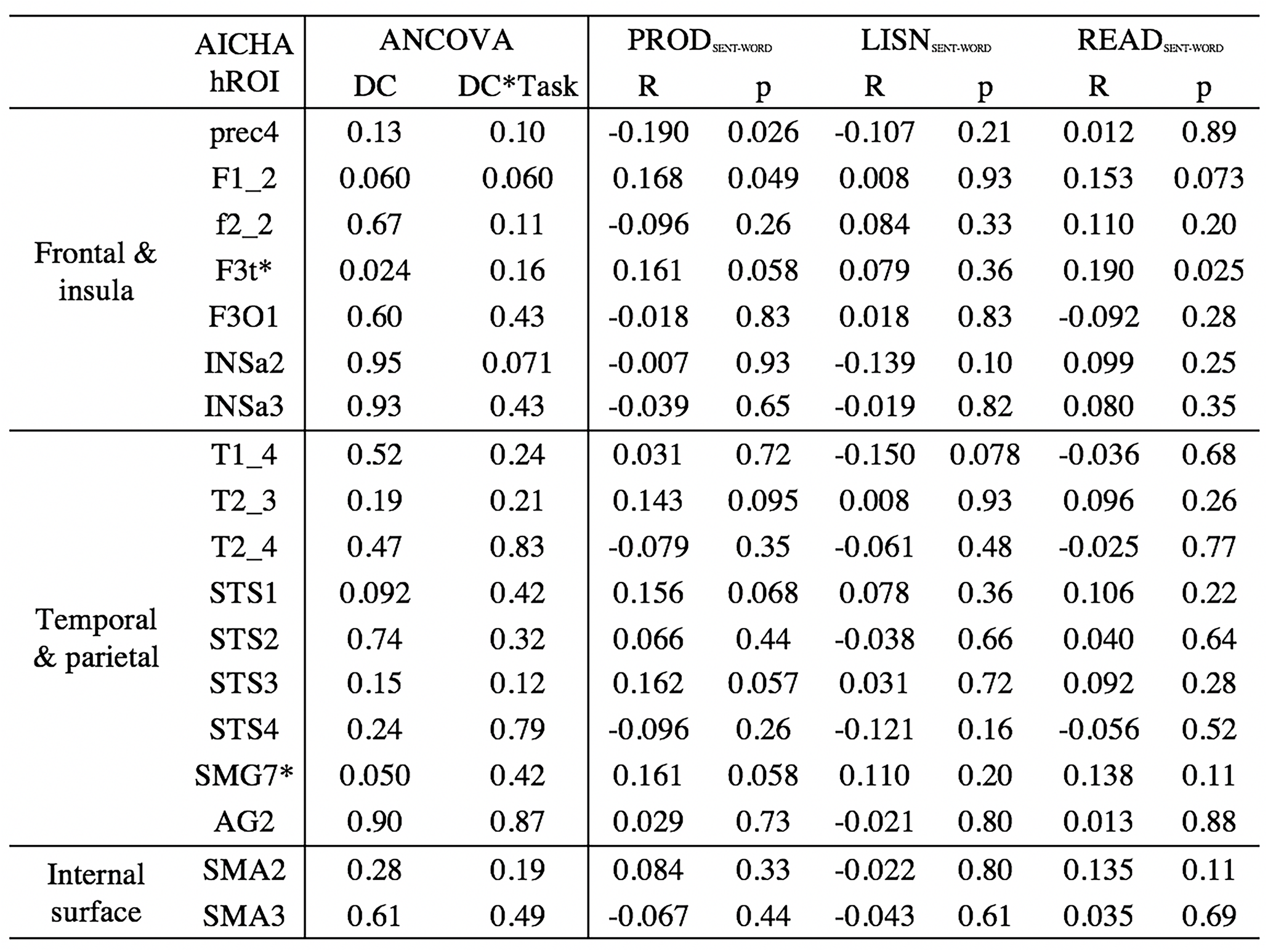
SENT_CORE hROIs correlations (R) between the degree centrality (DC) values calculated across the 185 hROIs of the left hemisphere and the mean activation in each of the 3 language tasks. hROIs with a star (*) are those having a significant correlation (p < 0.05) between the language tasks activation values and DC values.

